# The epigenetic modifier DOT1L regulates gene regulatory networks necessary for cardiac patterning and cardiomyocyte cell cycle withdrawal

**DOI:** 10.1101/2022.10.18.512679

**Authors:** Paola Cattaneo, Michael G. B. Hayes, Nina Baumgarten, Dennis Hecker, Sofia Peruzzo, Paolo Kunderfranco, Veronica Larcher, Lunfeng Zhang, Riccardo Contu, Gregory Fonseca, Simone Spinozzi, Ju Chen, Gianluigi Condorelli, Marcel H Schulz, Sven Heinz, Nuno Guimarães-Camboa, Sylvia M. Evans

## Abstract

Mechanisms by which specific histone modifications regulate distinct gene regulatory networks remain little understood. We investigated how H3K79me2, a modification catalyzed by DOT1L and previously considered a general transcriptional activation mark, regulates gene expression in mammalian cardiogenesis. Early embryonic cardiomyocyte ablation of *Dot1l* revealed that H3K79me2 does not act as a general transcriptional activator, but rather regulates highly specific gene regulatory networks at two critical cardiogenic junctures: left ventricle patterning and postnatal cardiomyocyte cell cycle withdrawal. Mechanistic analyses revealed that H3K79me2 in two distinct domains, gene bodies and regulatory elements, synergized to promote expression of genes activated by DOT1L. Surprisingly, these analyses also revealed that H3K79me2 in specific regulatory elements contributed to silencing genes usually not expressed in cardiomyocytes. As DOT1L mutants had increased numbers of postnatal mononuclear cardiomyocytes and prolonged cardiomyocyte cell cycle activity, controlled inhibition of DOT1L might be a strategy to promote cardiac regeneration post-injury.

## INTRODUCTION

Epigenetic enzymes play critical roles in organogenesis by defining chromatin structures required for cell type-specific transcriptional networks^1^. Yet, mechanisms by which genome-wide histone modifications result in activation or repression of specific genes remain little understood. DOT1L is the only epigenetic enzyme catalyzing methylation of lysine 79 of histone 3 (H3K79me)^2,3^. Originally identified in yeast due to its role in maintenance of telomeric regions^2^, DOT1L has been extensively studied in MLL-rearranged leukemia, where it is considered an emerging therapeutic target^4^. Initial genome-wide studies suggested that, rather than being associated with specific gene expression programs, H3K79me2/3 is ubiquitously found in all transcribed loci, leading to a model of H3K79me as a generic mark of active genes^5^.

Recent studies have revealed that, in addition to gene body H3K79me, DOT1L can regulate expression of its targets via methylation of regulatory regions^6^. We have previously shown that, *in vitro*, DOT1L is required for proper differentiation of embryonic stem cells into cardiomyocytes^7^, however it is currently unclear whether this enzyme is necessary for normal cardiogenesis *in vivo*. In mice, global ablation of *Dot1l* results in embryonic lethality^8^ and selective *Dot1l* ablation in cardiomyocytes results in an adult lethal phenotype with perturbations in dystrophin expression^9^. However, owing to the nature of this knockout (using a Cre that is not active during early embryogenesis), it is not known whether DOT1L plays any role in cardiogenesis. Furthermore, our molecular understanding of the functioning of this enzyme in cardiogenesis *in vivo* is limited by the absence of RNA-seq or ChIP-seq datasets that would allow for unbiased identification of direct targets of DOT1L in embryonic cardiomyocytes.

Cardiac patterning, the process allowing differentiation of the distinct cardiac chambers during embryogenesis, is controlled by the asymmetric expression of a well-defined set of transcription factors^10-13^. Amongst these, *Hand1* (expressed in the left ventricle) and *Hand2* (predominantly expressed in the right ventricle) play a major role in defining systemic and pulmonary ventricular identity, respectively^14-16^. Epigenetic mechanisms contributing to the tightly regulated expression of patterning genes remain mostly unexplored, however, it is known that the H3K4 methyltransferase SMYD1/BOP1 is a critical regulator of *Hand2* expression^17^. To date, it is not known whether any particular epigenetic enzyme plays a similar role in left ventricle-specific regulation of *Hand1* expression.

Regulated cardiomyocyte proliferation is a major component of normal cardiogenesis. The rate of cardiomyocyte mitosis gradually decreases from midgestational stages until birth, reaching a state of complete mitotic withdrawal in the first week after birth, when the majority of cardiomyocytes become binucleated^18^. Finding epigenetic enzymes that play a role in this process might allow the identification of routes to promote adult cardiomyocyte proliferation for therapeutic purposes.

To test whether DOT1L plays a role in cardiogenesis, we ablated a floxed *Dot1l* allele using a Cre allele that is active in cardiomyocytes from early developmental timepoints and observed a perinatal lethal phenotype with abnormal cardiac morphology. This phenotype resulted from perturbations in highly specific gene expression programs that drive two critical cardiogenic events: cardiac patterning and cardiomyocyte cell cycle withdrawal. In embryonic development, DOT1L emerged as a major regulator of the expression of several transcription factors involved in cardiac patterning, with left ventricle-specific genes being particularly sensitive to the absence of this enzyme. In the neonatal period, DOT1L promoted expression of genes involved in cardiomyocyte maturation and cell cycle exit. Integration of cardiomyocyte ChIP-seq and Hi-C data revealed that DOT1L regulated target genes via two mechanisms: methylation of H3K79 in gene bodies and methylation of H3K79 in regulatory elements (K79-REs). Our analyses identified two types of K79-REs: activating elements that synergized with gene body H3K79me2 to promote expression of targets activated by DOT1L, and K79-REs that contributed to silencing of genes normally not expressed in cardiomyocytes. In contrast to H3K79me2 being a general activator of gene transcription as previously thought, our results reveal H3K79me2 as an epigenetic mark that regulates specific gene programs by both activating and repressing target genes. Moreover, the increased proliferation of postnatal cardiomyocytes following loss of *Dot1l* suggests the use of DOT1L inhibitors, already in development for treatment of leukemia^19,20^, as agents to promote cardiac regeneration.

## RESULTS

### Abnormal cardiogenesis in DOT1L cKOs

Following our previous observation that DOT1L is required for the differentiation of embryonic stem cells into cardiomyocytes^7^, we decided to assess if this enzyme is required for normal cardiogenesis *in vivo*. Fluorescent in situ hybridization studies showed that the *Dot1l* gene is robustly expressed in cardiomyocytes at least from E10.5 onwards (Supplementary Figure 1a). This expression was observed throughout the heart, without being specific to any chamber. *Dot1l* expression was also observed in multiple tissues outside the heart, showing this enzyme is not cardiac-specific. However, at E16.5, *Dot1l* expression levels were higher in cardiomyocytes than in the developing valves (Supplementary Figure 1b). *Dot1l* has been previously ablated in the heart using an *αMHC-Cre* line, leading to an adult lethal phenotype with reduced dystrophin expression^9^. We have shown that *αMHC-Cre* is not optimal for myocardial-restricted knockouts due to its expression in non-myocyte lineages inside and outside the heart, and because it does not flox-out in all cardiomyocytes at an early time point^21^. To avoid these limitations and study cardiomyocyte-specific roles of DOT1L from early embryonic timepoints, we ablated a floxed *Dot1l* allele (loxP sites flanking exon 2) using the *xMlc2-Cre* allele^22^ that drives highly specific and efficient recombination in cardiomyocytes (Supplementary Figure 1c) from the cardiac crescent stage^22^. In our analyses, we compared *xMlc2-Cre*+; *Dot1l-WT/flox* mice (herein designated as Dot1L Ctrl or controls) with *xMlc2-Cre*+; *Dot1l-Δ/flox* littermate mice (herein designated as Dot1L cKO or mutants). Inclusion of a copy of the *Rosa26-tdTomato* reporter allele^23^ in all crosses allowed highly specific labeling of cardiomyocytes by the red fluorescent protein tdTomato for downstream flow cytometry and confocal microscopy applications (Supplementary Figure 1d). Real time qPCR analyses revealed highly efficient ablation of the floxed allele in sorted E12.5 cardiomyocytes (Figure 1a).

**Figure 1:**
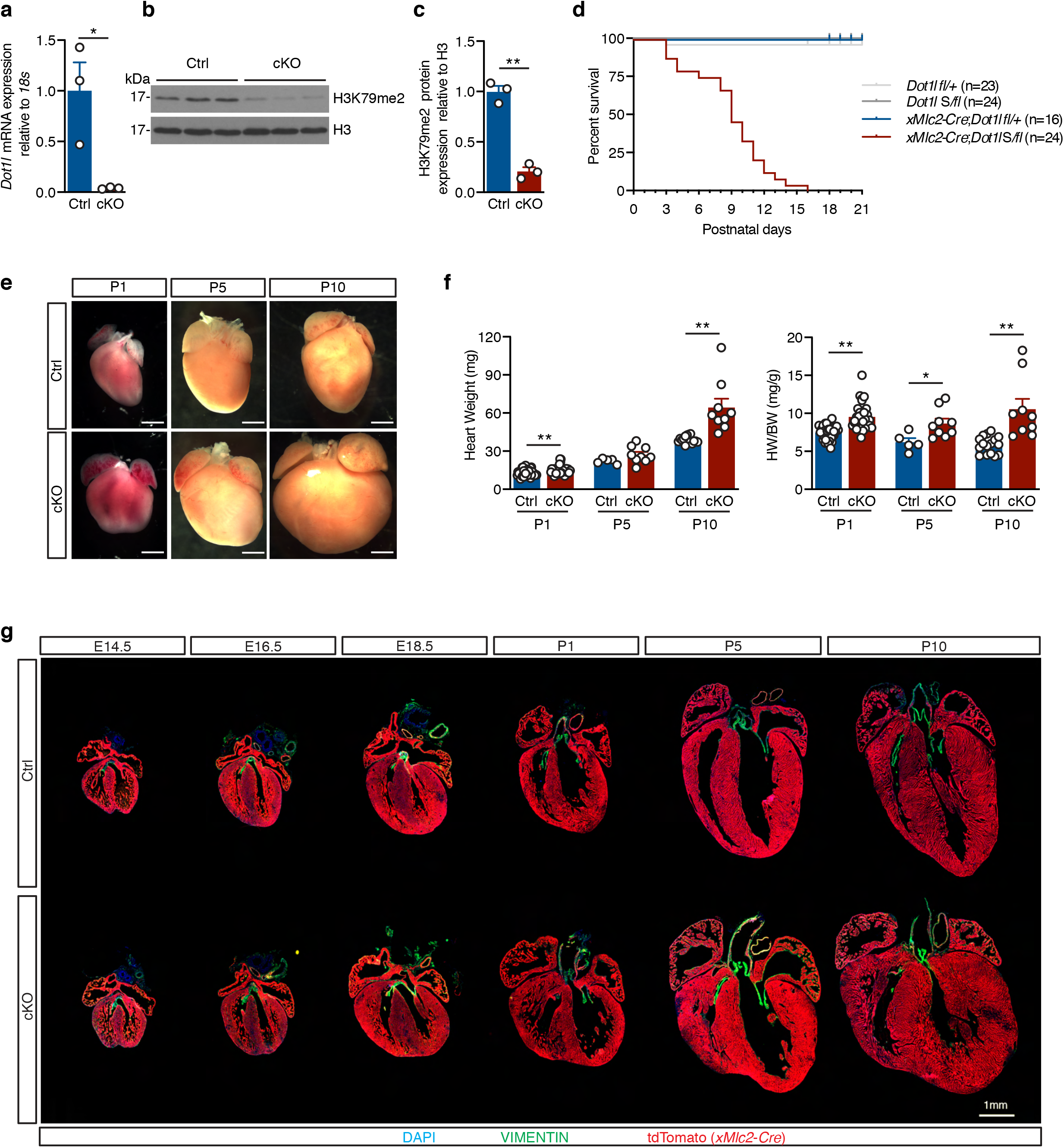
Cardiomyocyte-specific ablation of DOT1L from early developmental timepoints results in enlarged hearts and peri-natal lethality. **a)** qPCR analysis using a primer within the floxed exon of *Dot1l* mRNA showing efficient ablation of this gene in E12.5 cKO cardiomyocytes (N=3 biological replicates). **b-c)** Western blot (b) and respective quantification (c) showing strongly reduced H3K79me2 levels in E14.5 hearts upon ablation of Dot1L (N=3 biological replicates). **d)** Kaplan-Meier survival curves showing postnatal lethality of Dot1L cKOs. **e)** Whole mount images of postnatal day (P) 1, P5 and P10 hearts in Ctrls and Dot1L cKOs representative of the enlarged heart phenotype of Dot1L cKOs (scale bars = 1mm). **f)** Graphs representing a significant increase in heart weight (mg) and heart weight/body weight ratio (HW/BW (mg/g)) in Dot1L cKO vs Ctrl in all stages analyzed. **g)** Immunofluorescence time course depicting the dynamics of phenotypic manifestations in Dot1L cKOs in embryonic (E) and postnatal (P) stages. DAPI (4’,6-diamidin-2-phenylindol) in blue, VIMENTIN in green and lineage traced XMLC2-Cre;tdTomato cardiomyocytes in red (scale bar = 1mm). In all graphs Ctrl indicates control mice (*XMlc2-Cre;Dot1L fl/+)*, cKO indicates mutant mice (*XMlc2-Cre;Dot1L Δ/fl*). Data is presented as mean ± SEM; * P≤0.05, **P≤0.01.

DOT1L is the only enzyme catalyzing H3K79 methylation^3^. Consequently, cardiomyocyte ablation of *Dot1l* resulted in a strong reduction in H3K79me2 levels in E14.5 hearts (Figure 1b and 1c). Cardiomyocyte-specific ablation of *Dot1l* did not result in an embryonic lethal phenotype, as animals with the mutant genotype were observed at expected Mendelian ratios in all stages analyzed (Supplementary Figure 1e). Despite being born at expected numbers, Dot1L cKO mice started dying shortly after birth, with 50% mortality by postnatal day 9 (P9), and the longest-lived mutant surviving until P16 (Figure 1d). Macroscopic examination of organs at neonatal stages revealed that mutants had enlarged hearts with a rounded shape relative to controls (Figure 1e). This phenotype was exacerbated as animals aged and peaked at P10 in surviving animals (Figure 1e). This evident cardiac enlargement translated into increased heart weight and increased heart weight/body weight ratios (Figure 1f) without changes in body weight (Supplementary Figure 1f). To determine the timing of onset of this phenotype and assess for additional morphogenic malformations, we conducted a histological time course analysis spanning from midgestation to postnatal timepoints (Figure 1g). cKO hearts were completely indistinguishable from control littermates until E14.5 and started exhibiting the enlarged, rounded phenotype between E16.5 and E18.5. Postnatally, mutant hearts displayed ventricular walls with increased thickness and areas of moderate persistent trabeculation (Figure 1g). No additional major morphogenic abnormalities (septal or outflow defects) were observed.

Immunostaining for VIMENTIN revealed absence of valve malformations and absence of major foci of fibrosis in cKOs (Figure 1g).

### DOT1L is Essential for Cardiac Patterning

To decipher gene expression networks misregulated in the absence of DOT1L, we used RNA-seq to assess the transcriptomes of FACS-sorted Ctrl and cKO cardiomyocytes just prior to the onset of clear phenotypic changes (E16.5). An average of 100,000 cardiomyocytes were sorted per E16.5 heart (including atrial and ventricular chambers), yielding enough RNA to prepare biological replicates from individual hearts (3 replicates per genotype group).

Bioinformatics analyses revealed that from a total of 12110 expressed genes (RPKM ≥ 1 in either Ctrl or cKO), 1478 were misregulated in cKO versus Ctrl cardiomyocytes (log2FC ≤ −0.5 or ≥0.5, false discovery rate ≤0.05). From these, 439 genes were downregulated and 1039 upregulated in cKOs (Figure 2a and Supplementary File 1a). Transcripts upregulated in cKOs corresponded to genes that are normally expressed at low levels in control cardiomyocytes (62% of genes in the lower expression quartile), whereas downregulated genes corresponded to genes normally expressed at a medium/high level (Figure 2b). Notably, the gene displaying the highest downregulation (−5.4 log2FC) in mutant cardiomyocytes was the transcription factor *Hand1*, a central element in the establishment of left ventricular identity (Figure 2c and Supplementary File 1a). In addition, amongst genes significantly downregulated in mutants there were several other genes involved in cardiac patterning: the transcription factors *Irx4* (ventricular specific^24^), *Tbx5* (left ventricle and atrial-specific^25^), *Gata4 and Mef2c*, the epigenetic enzyme *Smyd1* and the left ventricular-specific genes *Gja5* (encoding Connexin40) and *Cited1* (Figure 2c and Supplementary File 1a). The transcription factor *Nkx2-5* also exhibited a trend toward downregulation (Supplementary File 1a). These transcriptomic differences revealed that DOT1L activity in embryonic cardiomyocytes is essential for establishment of gene expression networks that coordinate normal cardiogenesis. This requirement, however, was not generalized to all cardiac patterning genes, as transcript levels for *Hand2, Tbx20* and *Nppa* (ANF) were not altered in mutants (Supplementary File 1a).

**Figure 2:**
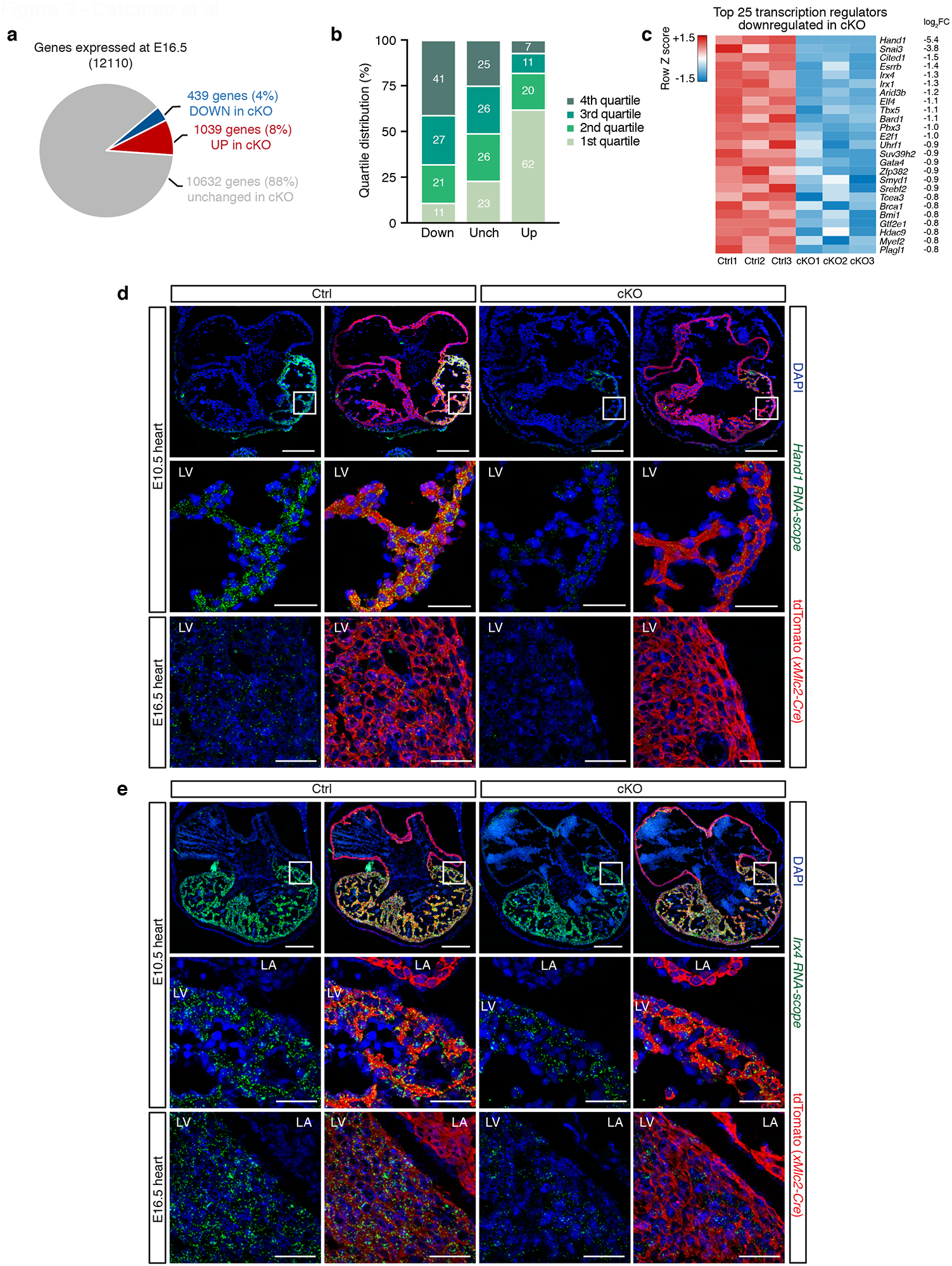
DOT1L is required in cardiomyocytes for normal expression of cardiac patterning genes. **a)** Pie chart representing the number of genes down-(log2FC ≥ −0.5; FDR ≤ 0.05) and up-regulated (log2FC ≥ 0.5; FDR ≤ 0.05) in Dot1L cKO cardiomyocytes at E16.5. **b)** Quartile distribution of gene expression in cardiomyocytes at E16.5. Genes downregulated (Down) in Dot1L cKO were expressed at a high level in control cardiomyocytes, whereas the majority of upregulated genes (Up) belonged to the bottom quartile of expression. Genes not significantly modulated (Unch) were evenly distributed across quartiles of expression. Data are shown as stacked percentage bar graph. **c)** Heatmap showing the expression of the top 25 transcription regulators downregulated in Dot1L cKO cardiomyocytes at E16.5, highlight that multiple transcription regulators with known roles in patterning of the cardiac chambers were significantly downregulated. **d-e)** RNA-scope analyses validating blunted expression of *Hand1* (d) and reduced levels of *Irx4* (e) in Dot1L cKOs both at E10.5 and E16.5. Scale bars = 250μm for low magnification panels and 50μm for high magnification.

To confirm the hypothesis of abnormal cardiac patterning in Dot1L cKOs, we performed fluorescent RNA in situ hybridization (RNA-scope) in histological sections of control and cKO hearts. Strikingly, at E16.5, *Hand1* transcripts were completely undetectable in the left ventricle of mutants (consistent with the 0 RPKM values observed in RNA-seq studies, (Figure 2d, bottom panel). Downregulation of *Hand1* was already evident at E10.5 when levels of *Hand1* expression in control hearts were higher than at E16.5 (Figure 2d, top and middle panels), indicating that transcriptional consequences of DOT1L ablation preceded the first phenotypic manifestations by several days. Similarly, transcript levels for the ventricular-specific transcription factor *Irx4* were significantly reduced in cKO hearts both at E10.5 and E16.5 (Figure 2e), confirming RNA-seq results.

Our data revealed a previously unrecognized role for DOT1L in regulating the expression of transcription factors involved in cardiac patterning, in particular those involved in left ventricular identity. While a number of genes were downregulated in cKOs, *Hand1* was particularly sensitive to absence of DOT1L, with *Hand1* transcripts being completely absent from midgestational DOT1L-deficient cardiomyocytes. Consistent with these transcriptomic observations, cardiac-specific ablation of *Hand1* phenocopies DOT1L cKO, including an enlarged and round-shaped heart and peri-natal lethality^16^.

### Transcriptional Control Via Gene Body H3K79me2

Several core cardiac transcription factors are differentially expressed in DOT1L cKO cardiomyocytes. Thus, genes modulated in our transcriptomic analyses likely include a combination of targets directly regulated by DOT1L as well as those indirectly regulated owing to secondary effects. To identify genes directly regulated by DOT1L, we performed ChIP-seq assays for H3K79me2 in E16.5 control and cKO FACS-sorted cardiomyocytes. These analyses revealed a total of 31,895 H3K79me2 peaks significantly enriched in control versus DOT1L cKO CMs (Supplementary File 2). The fact that the vast majority of H3K79me2 peaks in Ctrls were lost in cKOs is consistent with DOT1L being the sole histone K79 methyltransferase in these cells and validated the purity of our sorted CMs (Figure 3a and Supplementary File 2). Genome-wide differential peak distribution analysis revealed that H3K79me2 is mostly an intragenic modification (introns, exons, UTR and promoter/TSS) with only 2% of differential peaks being located in intergenic regions (Figure 3b).

**Figure 3:**
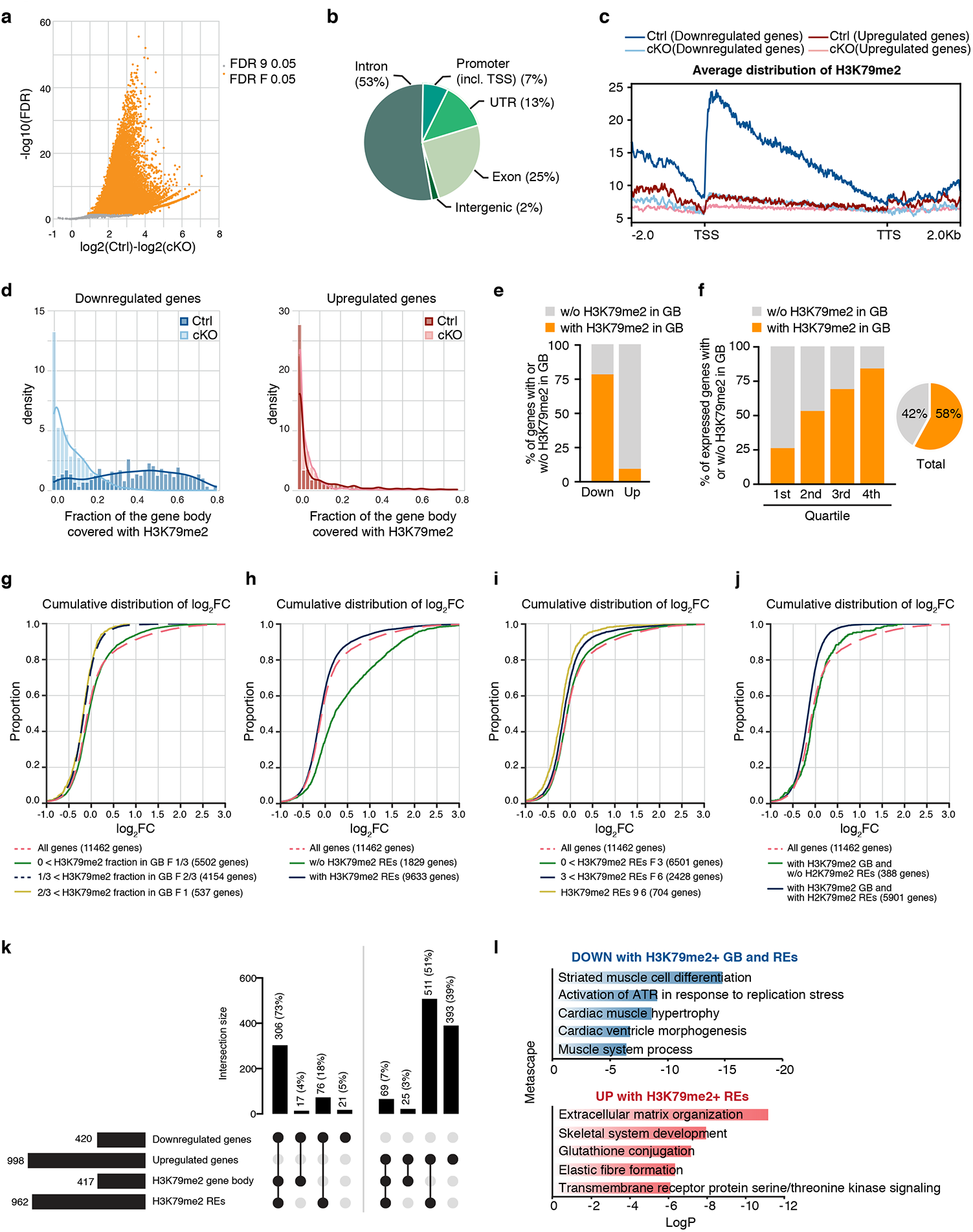
DOT1L controls transcription of target genes via a combination of H3K79me2 in gene bodies and regulatory elements. **a)** Volcano plot displaying H3K79me2 ChIP-seq peaks significantly enriched in E16.5 Dot1L Ctrl vs cKO cardiomyocytes. **b)** Pie chart indicating the genomic distribution of differential H3K79me2 ChIP-seq peaks in E16.5 cardiomyocytes. **c)** Metagene profiles showing the average distribution of H3K79me2 input-normalized tag density relative to Transcription Start Site (TSS) and Transcription Termination Site (TTS) with ±2Kb flanking regions. Genes downregulated in Dot1L cKOs had, in control cardiomyocytes, high levels of H3K79me2 in the vicinity of the TSS that progressively decreased towards the TTS. Genes upregulated in Dot1L cKOs did not show significant levels of H3K79me2 in Ctrl or cKO cardiomyocytes. **d)** Fraction of gene body covered with H3K79me2 in downregulated genes (left graph) and in upregulated genes (right graph) indicating that most genes downregulated in E16.5 Dot1L cKOs had, in Ctrl cardiomyocytes, H3K79me2 covering the majority of the gene body, whereas the majority of genes upregulated in cKOs didn’t have H3K79me2 gene body coverage in Ctrl cardiomyocytes. **e)** Graph representing the percentage of down- and up-regulated genes in E16.5 cKO cardiomyocytes with (Coverage ≥ 50 reads and Fraction of gene body ≥ 0.2) or without (Coverage < 50 reads and Fraction of gene body < 0.2) gene body H3K79me2 in E16.5 Ctrl cardiomyocytes. **f)** Percentage of genes with H3K79me2 in the gene body (GB) (Coverage ≥ 50 reads and Fraction of gene body≥ 0.2) or without H3K79me2 in the gene body (Coverage <50 reads and Fraction of gene body< 0.2). This modification was abundant amongst highly expressed genes (4^th^ quartile of RNA expression) and progressively decreased towards the lower quartiles of expression. Globally more than half (58%) of all genes expressed in E16.5 cardiomyocytes had gene body H3K79me2. **g-h-i-j)** Cumulative log2FC distribution analyses in different categories of genes in E16.5 cardiomyocytes: genes with different levels of H3K79me2 fraction in the gene body (GB) (g); genes with and without H3K79me2/H3K27ac positive regulatory elements (REs) (h); genes with increasing numbers of H3K79me2/H3K27ac positive regulatory elements (REs) (i); genes with a combination or none of H3K79me2 in GB and REs (j). DOT1L regulated expression of target genes via H3K79me2 in GB and REs and genes associated with both types of H3K79me2 (GB and RE) were more downregulated in cKOs than those containing exclusively gene body H3K79me2. A Kolmogorov-Smirnov test was applied to assess statistical significance of differences between distribution of gene groups. **k)** UpSet plot indicating the number and percentage of genes up and downregulated in cKOs vs Ctrls with and/or without H3K79me2 in gene body (GB) and regulatory elements (REs) in E16.5 cardiomyocytes. **l)** Metascape pathway analysis of genes downregulated with H3K79me2 in GB and REs (top) and upregulated with H3K79me2 in REs (bottom) in Dot1L cKO vs control E16.5 cardiomyocytes. Top 5 enriched categories are shown, sorted by Log P value.

Genes downregulated in cKOs exhibited, in control CMs, maximal levels of H3K79me2 immediately after the transcription start site (TSS) that progressively decreased towards the transcription termination site (TTS) (Figure 3c). Gene body H3K79me2 (coverage≥50 reads and fraction of gene body≥0.20 in control) was present in 341 downregulated genes (77% of all downregulated genes) and 98 upregulated genes (9% of all upregulated genes) (Figure 3d, 3e and Supplementary File 1). This observation suggests that a significant part of gene downregulation observed in cKOs can be directly attributed to H3K79me2 in the gene body, whereas gene upregulation does not seem to be directly associated with gene body H3K79me2. This is consistent with the view of gene body H3K79me2 as an activation mark^5^, however, it should be noted that this modification is not an absolute requirement for transcription, as multiple genes were expressed in controls without having significant levels of gene body H3K79me2 (Figure 3f). Within genes showing gene body H3K79me2 there were two clear categories: genes dependent on DOT1L (modulated in cKOs) and genes not significantly affected by the loss of DOT1L, suggesting a scenario in which other epigenetic modifications can compensate for the loss of gene body H3K79me2 in a subset of genes. Notably, the majority of genes whose transcription depended on gene body H3K79me2 belonged to the highest quartiles of expression, suggesting this modification might play a role in supporting high transcriptional levels.

### Transcriptional Control Via H3K79me2 in Regulatory Elements

Recent studies in leukemia showed that there is a distinct subset of enhancers that depend on H3K79me2^6^. We conducted bioinformatic analyses to investigate whether cardiomyocytes have H3K79me2-dependent regulatory elements (REs) that, together with gene body H3K79me2, contribute to gene expression differences observed in cKOs. To predict interactions between candidate regulatory elements and target genes, we applied the Activity-by-Contact (ABC) model^26^. By integrating information on chromatin state (ChIP-seq) and genomic conformation (Hi-C), this method is more accurate at predicting actual interactions between REs and their target genes than methods simply based on physical proximity^26^. Because the promoter of a gene can serve as a RE for a distinct cis gene^27,28^, our analyses also considered interactions between promoters and distal genes. We used the ABC algorithm to merge our own H3K79me2 ChIP-seq datasets with publicly available datasets of E16.5 heart H3K27ac ChIP-seq, for the identification of regulatory regions^29^ (ENCODE project, ENCFF153DSU), and of cardiomyocyte Hi-C^30^ (GSM2544836). To ensure we focused on meaningful RE-target gene interactions, in all downstream analyses we only considered the top 5% most relevant interactions identified by the ABC method: 471645 interactions, corresponding to 63958 H3K27ac peaks (Supplementary File 3). From these, 25% (15991 peaks) overlapped with a differential H3K79me2 peak (Supplementary File 3). This large number of cardiomyocyte REs enriched in H3K27ac and H3K79me2 (from here on designated as K79-REs) could provide an alternative mechanism for gene expression regulation by DOT1L. To assess the relative contribution of H3K79me2 in gene body versus H3K79me2 in REs to the regulation of target genes, we quantified the effect of these variables in the cumulative distribution of log2FC values. These analyses revealed that genes with higher fraction of H3K79me2 in the gene body were more downregulated in mutants than genes with lower (less than 33%) fraction of H3K79me2 in the gene body (Figure 3g and Supplementary File 4). Considering all genes, those that interacted with K79-REs displayed a trend to being downregulated in mutant cardiomyocytes, whereas genes that did not interact with these REs were more likely to be upregulated (Figure 3h and Supplementary File 4). The overall contribution of K79-REs to the relative expression of target genes seemed less evident than the effect of gene body H3K79me2 (Figure 3g,h). However, a stratified analysis segregating genes according to the number of associated K79-REs revealed that the magnitude of downregulation was relatively low if the gene had 1 to 3 K79-REs, but increased if the gene had 4 or more K79-REs (Figure 3i and Supplementary File 4). Focusing exclusively on loci with gene body H3K79me2, those that interacted with K79-REs were more downregulated in mutant cardiomyocytes than those without these elements, indicating that gene body and regulatory element H3K79me2 synergized to potentiate expression of target genes (Figure 3j and Supplementary File 4).

The vast majority of the genes downregulated in Dot1L-deficient cardiomyocytes were associated with both gene body H3K79me2 and at least one K79-RE (Figure 3k). Interestingly, 18% of all downregulated genes were associated with K79-REs in the absence of gene body H3K79me2. Overall, only 5% of downregulated genes were not associated with H3K79me2 in gene body or regulatory elements, indicating most gene downregulation was a direct consequence of DOT1L cKO. On the other side of the scale, upregulated genes had reduced association with gene body H3K79me2 (as mentioned above), but, surprisingly, 51% of upregulated genes were associated with K79-REs in the absence of gene body H3K79me2. This observation suggests a novel role for DOT1L in gene silencing via H3K79me2-dependent regulatory elements.

Functional annotation revealed that genes downregulated in cKO with H3K79me2 in gene bodies and REs were involved in muscle differentiation and morphogenesis (Figure 3l). On the other hand, genes upregulated in cKOs and associated with inhibitory K79-REs were related to non-myocyte functions: extracellular matrix organization and skeletal system development. To search for cues as to how DOT1L achieves both gene activation and silencing via methylation of REs, we screened for enrichment in binding sites for known transcriptional regulators in the K79-REs of genes downregulated versus genes upregulated in DOT1L cKOs. This analysis revealed that K79-REs associated with genes downregulated in DOT1L cKOs (activating K79-REs) were enriched in binding sites for TFs of the Forkhead box (FOX) family, known regulators of multiple steps in mammalian cardiogenesis^31^, as well as multiple cardiac-specific TFs (Figure 4c). On the other side of the scale, the most enriched element in K79-REs associated with genes upregulated in cKOs (silencing K79-REs) corresponded to the binding site of MBD2, a member of the NuRD complex known to play an important role in transcriptional silencing^32^ (Figure 4d).

**Figure 4:**
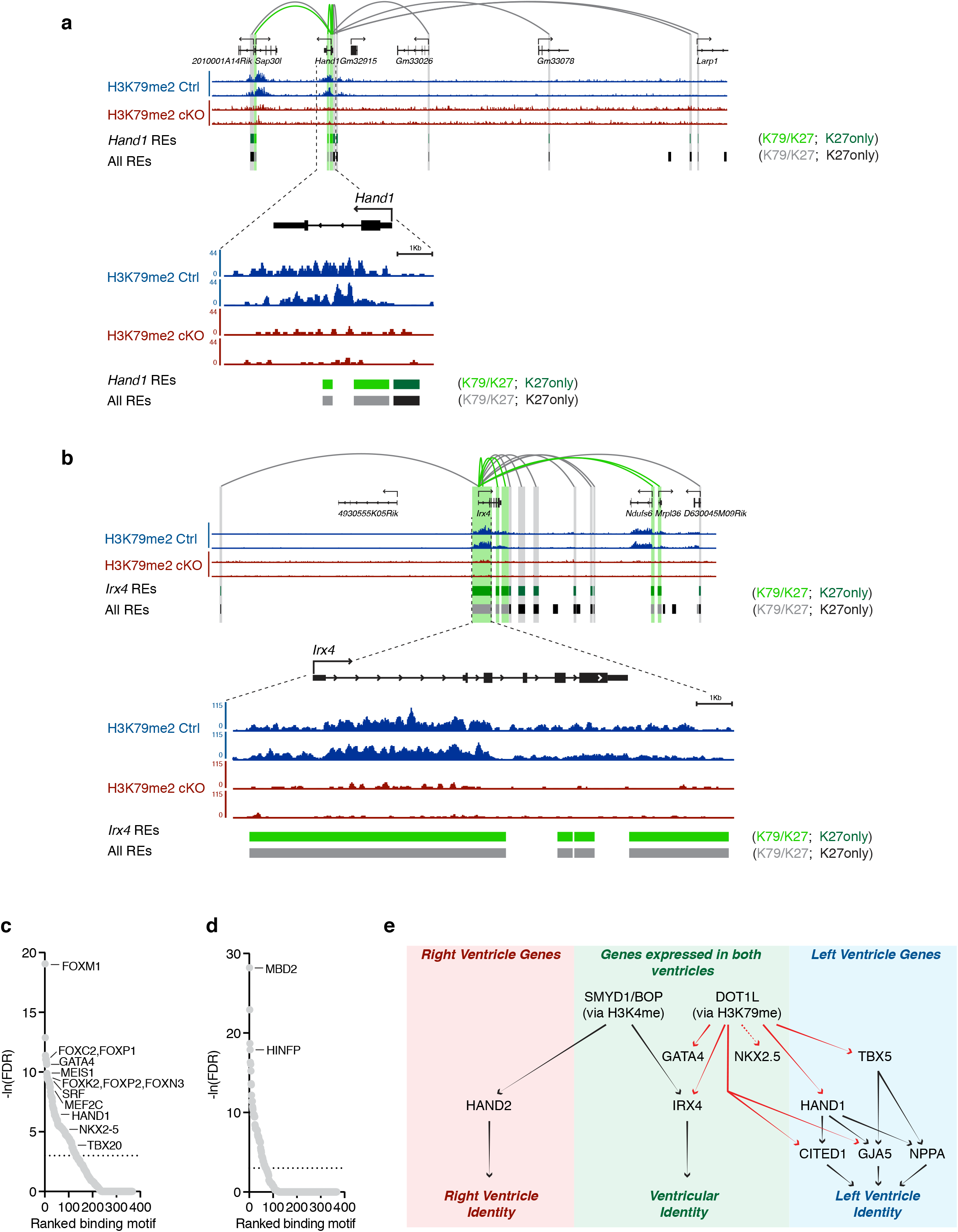
Genomic interactions underlying H3K79me2-dependent regulation of *Hand1* and *Irx4* transcription. **a, b)** Browser tracks displaying H3K79me2 ChIP-seq profiles of Ctrls (blue) and Dot1L cKOs (red) E16.5 cardiomyocytes in the genomic regions containing the *Hand1* (a) and *Irx4* (b) loci. Loops display all significant regulatory interactions between REs and the *Hand1* or *Irx4* loci, as identified by the AbC analysis. Gray loops identify interactions with REs that have H3K27ac but not H3K79me2, whereas green loops identify interactions with REs enriched in both epigenetic marks. For reference, all REs (defined as H3K27ac+ peaks) are displayed in the bottom lane, regardless of their regulatory association with the *Hand1* and *Irx4* loci. **c, d)** Motif enrichment analysis ranking transcription factors (TFs) with TRAP values enriched in REs associated with downregulated genes (c) and upregulated genes (d). **e)** Diagrammatic representation of key cardiac patterning genes directly (red arrows) or indirectly (black arrows) regulated by DOT1L (via H3K79me2).

Focusing on the cardiac patterning genes downregulated in DOT1L cKOs it became evident that these were directly targeted by DOT1L, with the majority belonging to the group of loci that had gene body H3K79me2 combined with K79-REs. This category included not only the most downregulated gene, *Hand1*, but also the other aforementioned central players in cardiogenesis: *Irx4, Gata4, Tbx5*, and *Mef2c* (Figure 4a and 4b and Supplementary File 4).

Interestingly, *Smyd1*, encoding the H3K4 methyltransferase necessary for right ventricular expression of *Hand2*^*17*^, was also a direct target of DOT1L, unveiling complex epigenetic mechanisms for regulation of cardiac patterning. Our results suggest DOT1L has particular importance in left ventricle patterning, as it regulates not only transcription factors specifying the identity of this chamber (*Hand1* and *Tbx5*), but also their direct downstream targets *Cited1* and *Gja5* (Cx40)^16^. On the other hand, *Nppa* (ANF), another classic target of both HAND1^16^ and TBX5^33^ was not altered in DOT1L cKOs, further highlighting the specificity and complexity of this epigenetic control (Figure 4e).

### Impaired cardiomyocyte cell cycle withdrawal in Dot1L mutants

Defective cardiac patterning can account for the abnormal morphology of Dot1L cKO hearts, however, it does not explain the increased wall thickness observed in postnatal mutant hearts (Figure 1g). As increased myocardial wall thickening coincides with the developmental period in which cardiomyocytes withdraw from cell cycle^18^, we hypothesized this process might be affected in Dot1L cKOs. Flow cytometry analyses of cells isolated from P1 hearts revealed that, at this stage, about 50% of all cardiac cells were cardiomyocytes (labeled by expression of the reporter gene tdTomato, Figure 5a). Assessment of EdU incorporation rates (24 hours EdU pulse) revealed that, in control hearts, 19% of all cardiomyocytes were proliferative, whereas this number increased to 26% in mutant hearts (Figure 5a), representing a statistically significant increase in cardiomyocyte proliferation (Figure 5b). In mice, by P10 most cardiomyocytes are withdrawn from cell cycle and 80% of cardiomyocytes are binucleated^18^. Consistently, fluorescent microscopy analyses of cardiomyocytes isolated from P10 hearts revealed that less than 3% of control cardiomyocytes were EdU+ (24 hours EdU pulse, Figure 5c and 5d), compared to 10% in mutant hearts (Figure 5c and 5d). Nucleation analysis revealed that DOT1L cKO hearts had almost twice as many mononucleated cardiomyocytes (34% in cKOs versus 19% in controls) at the expense of binucleated cardiomyocytes (62% in cKOs versus 80% in Ctrls) (Figure 5e). Quantification of proliferation ratios across different nucleation categories (Figure 5f) revealed that Dot1L cKOs had a moderate increase in the percentage of EdU+ mononucleated cardiomyocytes (8.1% in cKOs versus 5.2% in Ctrls) and a significant 4.9-fold increase in the percentage of EdU+ binucleated cardiomyocytes (9.1% in cKOs versus 1.86% in controls).

**Figure 5:**
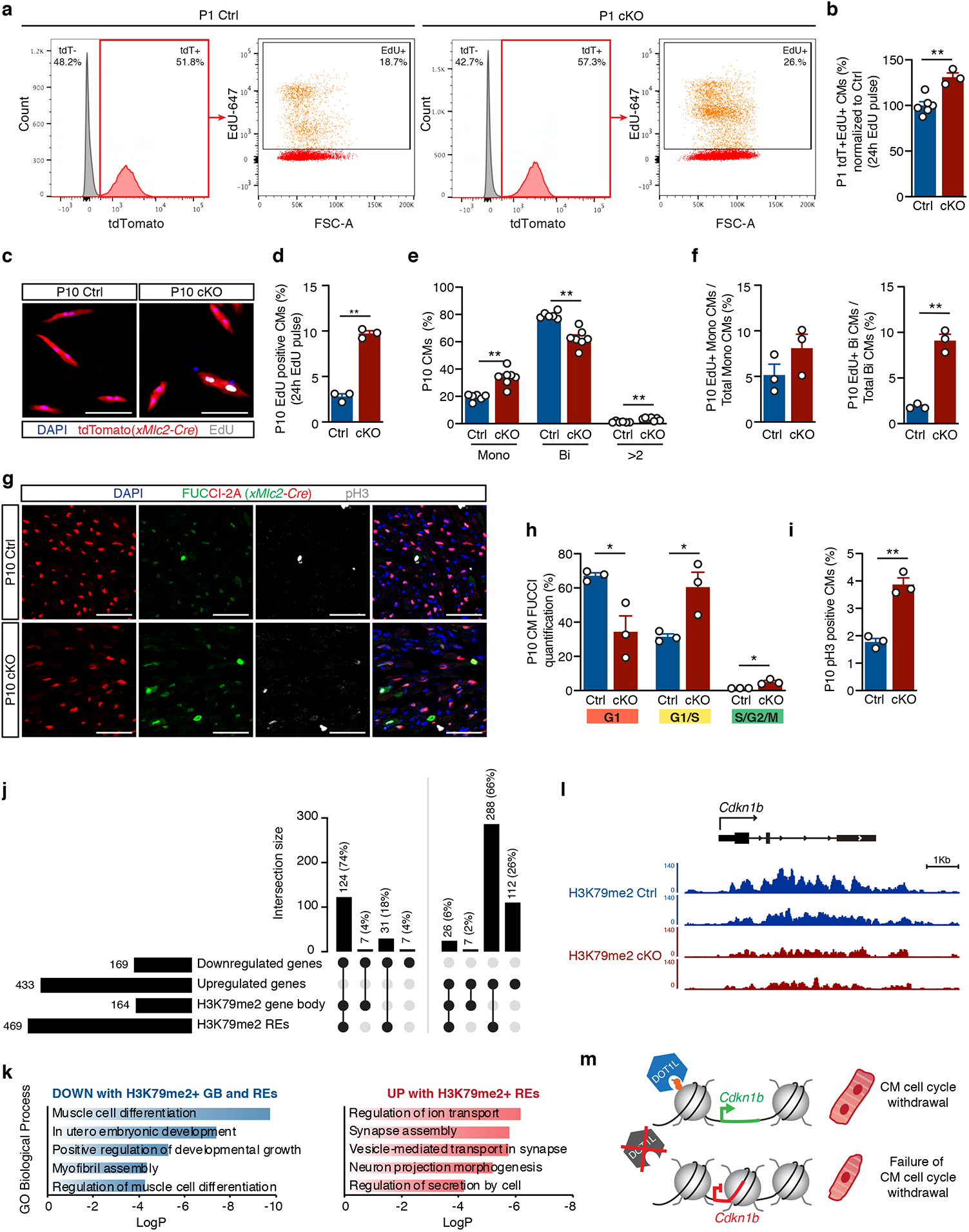
Neonatal Dot1L cKO cardiomyocytes fail to undergo cell cycle withdrawal. **a-b)** Representative FACS analysis (a) and respective quantification (b) showing significantly increased EdU incorporation within P1 cardiomyocytes (tdTomato+ cells) of Dot1L cKO vs Ctrl hearts (N:23 biological replicates). **c-d)** Representative immunofluorescence images (c), and respective quantification (d) showing significantly increased rates of EdU incorporation in P10 cardiomyocytes isolated from Dot1L cKO vs Ctrl hearts. DAPI in blue, endogenous tdTomato signal driven by XMlc2-Cre;Rosa26-tdTomato in red and EdU in white. (Scale bar =100μm; Mean of 755 CMs counted per heart from N=3 biological replicates). **e)** Quantification of relative percentage of mononuleated (Mono), binuclated (Bi) or multinucleated (>2) cardiomyocytes in P10 Dot1L Ctrls and cKOs. At P10, Dot1L cKO hearts had more mononucleated (Mono) and less binucleated (Bi) cardiomyocytes than littermate Ctrls (mean of 697 CMs counted per heart from N:26 biological replicates). **f)** Quantification of percentage of EdU+ cardiomyocytes within mononucleated (left graph) and binucleated (right graph) cardiomyocytes of P10 Dot1L Ctrls and cKOs (Mean of 755 CMs counted per heart from N=3 biological replicates). **g**,**h**,**i)** Immunofluorescence images (g) and respective quantification (h,i) of P10 Dot1L Ctrl and cKO hearts on a *Rosa26-*FUCCI2A background (Cre-dependent cell cycle indicator). Red-only nuclei represent cardiomyocytes in G1; red+green (yellow) nuclei represent cardiomyocytes in G1/S; green-only nuclei correspond to cardiomyocytes in S/G2/M, pH3 staining in white. Dot1L cKO hearts had a significantly higher percentage of cardiomyocytes in G1/S and in S/G2/M compared to Ctrls, (h, mean of 2070 CMs counted per heart from N=3 biological replicates) and of phophoHistone3+ (pH3) cardiomyocytes (i, mean of 1019 CMs counted per heart from N=3 biological replicates). (Scale bar = 50μm; sections have been quantified from all the compartment of the heart). **j)** UpSet plot indicating the number and percentage of genes up and downregulated in cKOs vs Ctrls with and/or without H3K79me2 in gene body (GB) and regulatory elements (REs) in P1 cardiomyocytes. **k)** Gene Ontology pathway analysis of genes downregulated with H3K79me2 in GB and REs (left) and upregulated with H3K79me2 in REs (right) in Dot1L cKO vs Ctrl P1 cardiomyocytes. Top 5 enriched categories are shown, sorted by Log P value. **l)** Browser tracks of H3K79me2 ChIP-seq in Ctrls (blue) and Dot1L cKOs (red) P1 cardiomyocytes in the genomic regions in the vicinity of *Cdkn1b* locus. **m)** Diagrammatic representation of mechanism of defective cardiomyocyte cell cycle withdrawal in the absence of DOT1L. In all graphs from (b) to (i) data is presented as mean ± SEM; * P≤0.05, **P≤0.01.

**Figure 6.**
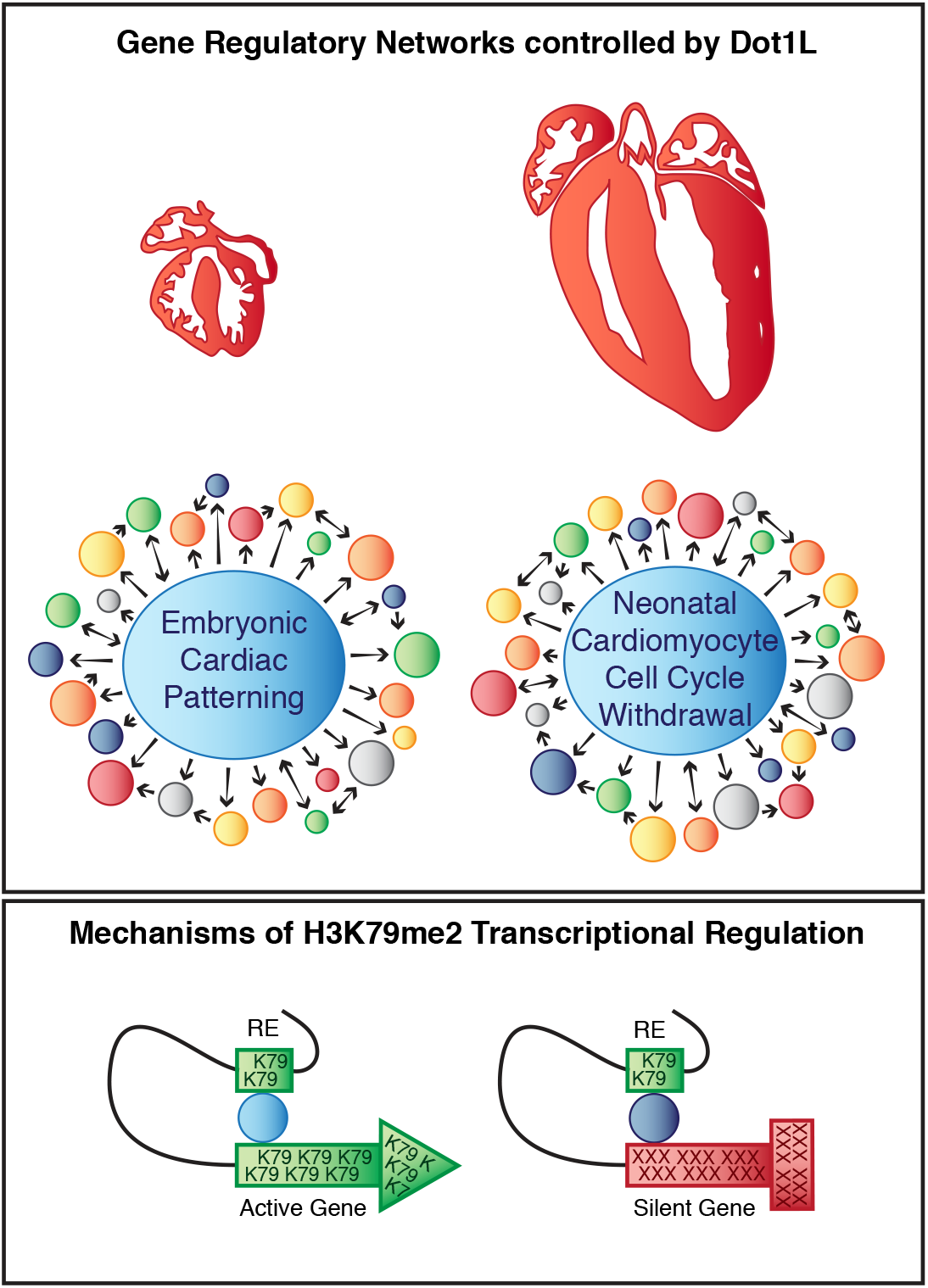
Graphical model of gene regulatory networks in embryonic and postnatal hearts controlled by Dot1L (top). Mechanisms of H3K79me2 transcription regulation (bottom).

To validate these analyses, we assessed proliferation of control and Dot1L cKO cardiomyocytes using the *Rosa26-Fucci2a* cell cycle reporter^34^. Expression from this allele is Cre-dependent (thus, restricted to cardiomyocytes in our analyses) and labels G1 cardiomyocytes in red, and actively proliferating cells in yellow (G1/S) or green (S/G2/M)^34^. Analysis of histological sections of P10 hearts (Figure 5g, 5h and Supplementary Figure 2a and 2d) revealed that in controls the majority of cardiomyocytes (67%) were in G1. G1/S cardiomyocytes could also be detected (32%), but S/G2/M cells were extremely rare (1%). On the other hand, DOT1L mutant cardiomyocytes showed significantly different cell cycle distribution across all cycling phases: 34% in G1, 61% in G1/S and 5% in S/G2/M (Figure 5g, 5h and Supplementary Figure 2a and 2d). In addition, increased proliferation of mutant cardiomyocytes was also confirmed by phospho-Histone 3 (pH3) antibody staining in histological sections (Figure 5g and 5i and Supplementary Figure 2b and 2c), as well as EdU analysis (2 hours EdU pulse, Supplementary Figure 2d-g) of Dot1L mutants and controls in the *Rosa26-Fucci2a* background.

To gain insight into gene expression networks underlying the sustained proliferation of Dot1L cKO cardiomyocytes, we used neonatal (P1) FACS-sorted cardiomyocytes to perform mechanistic analyses similar to those described for E16.5: RNA-seq, H3K79me2 ChIP-seq, followed by bioinformatic analysis employing the ABC method to predict interactions between regulatory elements and their target genes (Supplementary Figure 3 and Supplementary Files 1-4). These analyses revealed that, similar to E16.5, the majority of genes downregulated in cKO cardiomyocytes were directly regulated by DOT1L via H3K79me2 both in gene body and REs (Figure 5j). On the other hand, the vast majority of genes upregulated in cKOs (92%) did not have gene body H3K79me2, however, 66% had interactions with K79-REs (Figure 5j).

Functional annotation revealed that genes upregulated in cKOs and associated with K79-REs were enriched in genes involved in neuronal differentiation and vesicle-mediated transport in synapse, whereas genes downregulated in cKOs with H3K79me2 in gene body and REs were enriched in genes involved in muscle cell differentiation and myofibril organization (Figure 5k). Importantly, within this category of genes there was the gene encoding the cell cycle regulator p27 (*Cdkn1b*) (Figure 5l). Interestingly, p27 knockouts have a phenotype of increased heart size due to defective cardiomyocyte cell cycle arrest that resembles the one of Dot1L cKOs^35^.

Altogether these observations suggest that in the neonatal period, DOT1L directly promotes expression of genes involved in cardiomyocyte maturation and cell cycle withdrawal whilst repressing expression of genes associated with neuronal functions.

## DISCUSSION

Devising a blueprint for transcriptional and epigenetic mechanisms regulating normal heart development is of critical importance for our understanding of congenital heart disease and to pave the way for regenerative therapies of the adult heart post injury. Our previous studies using embryonic stem cells led to the hypothesis that DOT1L might play a relevant role in cardiogenesis, which we confirmed with the *in vivo* experiments reported here. In addition, we deciphered novel mechanisms of action of this epigenetic modifier. *Dot1l* is ubiquitously expressed in the heart and its cardiac-specific conditional ablation resulted in a fully penetrant phenotype of an enlarged and rounded heart that culminated in neonatal lethality (Figure 1).

Genome-wide transcriptomic and ChIP-seq assays provided detailed mechanistic insight into gene expression programs underlying this phenotype, highlighting DOT1L as a direct regulator of several genes involved in two processes that take place at distinct developmental stages: embryonic cardiac patterning (Figures 2, 3 and 4) and neonatal CM maturation and cell cycle withdrawal (Figures 5). Detailed bioinformatic analyses revealed that DOT1L regulated expression of its targets via methylation of H3K79 in gene body, regulatory regions, or a combination of both.

In embryonic cardiac patterning, DOT1L directly regulated several genes involved in defining distinct cardiac chambers. Amongst these, there was a clear enrichment in genes required for left ventricular identity (*Hand1, Tbx5, Cited1, Gja5*). From these, *Hand1* emerged as a gene particularly sensitive to absence of DOT1L activity. Likely reflecting this tight regulation, the dilated and rounded heart phenotype with neonatal lethality displayed by *Dot1l* mutants strongly resembles the phenotype of *Hand1* conditional mutants^16^. *Hand1* cardiac cKOs also display ventricular septal and outflow defects, whereas *Dot1l* mutants do not. This is likely a consequence of residual *Hand1* transcripts at earlier embryonic stages – *Hand1* transcripts were completely absent from E16.5 DOT1L cKO cardiomyocytes but could be detected at E10.5 (although at much lower levels than in stage-matched controls, Figure 2). The dependence of *Hand1* expression on DOT1L is of particular relevance for our understanding of epigenetic regulation of cardiogenesis. Previous studies have shown that SMYD1 (that catalyzes H3K4 methylation) is critical for activation of *Hand2*^17^, but an epigenetic mechanism specifically regulating *Hand1* has remained elusive. Together with these previous observations, our results suggest a model in which DOT1L and SMYD1, two enzymes expressed throughout the heart, are essential for left/right asymmetric gene expression by serving as regulators of critical patterning transcription factors (Figure 4c).

DOT1L depletion resulted in absence of *Hand1* expression without affecting expression of *Hand2*, whereas absence of SMYD1 results in blunted activation of *Hand2* without affecting expression of *Hand1*^*17*^. DOT1L also regulated expression of *Smyd1* itself, revealing a complex epigenetic mechanism controlling normal cardiogenesis. It is unclear if SMYD1 is involved in regulating *Dot1l* expression. Interestingly, despite the fact that *Smyd1* transcripts were significantly downregulated in Dot1L cKOs, its critical target *Hand2* was not altered in these mutants, suggesting that the levels of SMYD1 protein present in Dot1L cKOs are sufficient to sustain mechanisms of transcriptional regulation dependent on this enzyme. Despite divergent roles in the regulation of left/right genes, DOT1L and SMYD1 also share some common targets. For example, transcription of *Irx4* is dependent on the action of both enzymes (our own data and ^17^). IRX4 is required for ventricular myocyte identity, and it is conceivable that DOT1L and SMYD1 act in a coordinated way to ensure a rheostatic control of *Irx4* transcription. How DOT1L and SMYD1, two enzymes that are expressed throughout the heart, achieve regulation of chamber-specific genes (ventricular versus atrial, left versus right) is not currently known and will be the subject of future studies. Given that several of DOT1L targets are also targeted by critical regulators of cardiogenesis, such as NKX2.5 and TBX5, it is possible that the specificity of these epigenetic enzymes is derived from their interactions with distinct transcription factors.

Initial studies suggested that H3K79 methylation is a generic marker of active transcription^5^, but our datasets clearly revealed that, in cardiomyocytes, H3K79me2 methylation is not a basic requirement for gene transcription, as in control animals, multiple genes are active without having gene body H3K79 methylation. From the group of genes that bear this modification in controls and lose it in mutant hearts, two major categories emerge: those that are modulated upon DOT1L loss and those that are unaffected by the absence of this enzyme. The latter may reflect mechanisms of epigenetic redundancy. Our bioinformatic analyses integrating differential H3K79me2 ChIP-seq peaks with a mark of regulatory regions^29^ (H3K27ac ChIP-seq) and 3-dimensional genomic conformation (cardiomyocyte Hi-C) suggested a model in which, in addition to gene body H3K79me2, DOT1L mediates expression of target genes via H3K79me2 in cis-regulatory elements. In loci with gene body H3K79me2, those associated with K79-REs were more downregulated in Dot1L cKOs. The magnitude of downregulation correlated with two variables: extent of H3K79me2 gene body coverage and the number of K79-REs interacting with the gene, suggesting these two forms of H3K79me2 synergize to promote expression of target genes. Interestingly, our analyses suggested that K79-REs could also be of an inhibitory nature, blocking transcription of target genes that lack gene body H3K79me2. To our knowledge this is the first report of an involvement of DOT1L/H3K79me2 in this form of gene expression regulation.

In mice, after the first week of life, most cardiomyocytes are binucleated and completely withdrawn from the cell cycle^18^. Resistance of mature cardiomyocyte to proliferation is a major hurdle for cardiac regeneration^36,37^. Thus, devising strategies to promote reacquisition of proliferative potential in cardiomyocytes is a major priority in cardiac regenerative medicine^36,37^. FACS and histological studies using biochemical and genetic strategies to identify cardiomyocyte-specific proliferation revealed that Dot1L cKO cardiomyocytes retain proliferative potential at P10, a stage at which control counterparts are already withdrawn from the cell cycle (Figure 5). Notably, Dot1L cKO hearts also had a higher percentage of mononucleated cardiomyocytes than control hearts. Transcriptome analyses suggested that DOT1L depleted cardiomyocytes are less mature than control cardiomyocytes and fail to activate expression of p27, a known repressor of cell cycle. Permanent ablation of DOT1L results in severe consequences and lethality, but our results revealing sustained proliferation of DOT1L-depleted cardiomyocytes suggest that temporary inhibition of DOT1L in postnatal heart might be a strategy to promote re-acquisition of mitotic potential in cardiomyocytes. This prospect is particularly exciting considering that pharmacological inhibitors of DOT1L have been developed and are currently in clinical trials for leukemia^19,20^, and will be explored in follow up studies.

## METHODS

### Mouse Strains and Experiments

Animal experiments were conducted according to protocols approved by the Institutional Animal Care and Use Committee at University of California, San Diego. All transgenic lines used were kept on an outbred background (Black-Swiss, Charles River laboratories). Mice were maintained in plastic cages with filtered air intake ports (Techniplast) on a 12 hr light cycle and fed Teklad LM-485 irradiated diet (Harlan Laboratories, catalog number 7912). For analyses conducted in embryonic stages, embryos were staged according to the embryonic day (E) on which dissection took place, with noon of the vaginal plug day being considered as E0.5 and birth typically occurring at E19. *Dot1l*^flox^ mice were obtained from the KOMP Repository (CSD29070) https://www.komp.org/geneinfo.php?geneid=54455. Floxed-out *Dot1l*^*Δ*^ mice were generated by crossing the *Dot1l*^flox^ allele with the epiblastic *Meox2-Cre* allele obtained from JAX laboratories (Stock No: 026858). *xMlc2-Cre* mice^22^ were gently provided by Timothy Mohun. The *Rosa26-tdTomato (Ai9) (tdTom*) indicator allele^23^ was purchased from JAX (Stock No: 007905). The Rosa26-*Fucci2A* cell cycle reporter allele was gently provided by Ian James Jackson^34^. All experiments were performed using littermate cKOs and Ctrls. Images shown are representative examples of experiments with n≥3 biological replicates. In experiments assessing proliferation, mice received an injection of EdU (at P1 25μL of a 3g/L stock and at P10 50μL of a 3g/L stock) 2h or 24h prior to euthanasia.

For genotyping, genomic DNA was extracted by adding 250μL of 50mM NaOH to a tail tip biopsy and heating at 95°C for 30 minutes. The solution was then neutralized by adding 50μL of 1M Trish-HCl (pH 8.0). Genotyping PCR reactions (36 amplification cycles) were performed using Taq DNA Polymerase with ThermoPol Buffer (New England Biolabs, M0267L), dNTPs from Promega (U1511) and 1μL of DNA solution. The following genotyping primers were used:

XMlc2Cre-Fw 5’-TAGGATGCTGAGAATCAAAATGT-3’;

XMlc2Cre-Rev 5’-TCCCTGAACATGTCCATCAGGTTC-3’;

Dot1L-Fw 5’-CCATATTAGTGTTCAAGGGCTACT-3’;

Dot1L-fl/wt-Rev 5’-AGCATAAGGATGCCAACTACTAAC;

Dot1L-null-Rev 5’-AAGGAGGTCCTACTCATAGTCCTT-3’;

Rosa26-tdTomato-Fw 5’-CTGTTCCTGTACGGCATGG-3’;

Rosa26-tdTomato-Rev 5’-GGCATTAAAGCAGCGTATCC-3’;

Rosa26-wt-Fw 5’-AAGGGAGCTGCAGTGGAGTA-3’;

Rosa26-wt-Rev 5’-CCGAAAATCTGTGGGAAGTC-3’;

Rosa26-wt-Fw 5’-CAAAGTCGCTCTGAGTTGTTATCAG-3’;

Rosa26-wt-Rev 5’-GGAGCGGGAGAAATGGATATGAAG-3’;

Rosa26-Fucci2a-Rev 5’-TCACCCAGGAGTCATTTGAT-3’.

### Immunofluorescence

Tissues were isolated in cold PBS and fixed in 4% paraformaldehyde at 4°C overnight. Tissues were dehydrated in a sucrose gradient (5% −1 hour, 12% - 1 hour, 20% - overnight) and embedded in a 1:1 mix of 20% Sucrose and Tissue-Tek Optimal Cutting Temperature compound (OCT, Sakura). 10 μm thick histological sections were cut using a cryostat (Leica CM3050) and kept at -20°C (short-term) or −80°C (long-term) until being processed for immunofluorescence. Sections or isolated cardiomyocytes were blocked in 10% donkey serum before incubation with antibodies. Primary antibodies were incubated at 4°C overnight. The following primary antibodies were used: anti-α Sarcomeric Actinin (1:400 Abcam #ab68167), anti-Vimentin (1:100 Abcam#ab45939), anti-TNNT (1:100 Thermo Fisher #MA5-12960), anti-PDGFR-α (1:200 R&D Systems #AF1062), anti-tdTomato (1:100 Sicgen #ab8181-200), anti-GFP (1:600 Abcam #ab13970), anti-phosphoH3 (1:100 Millipore #06-570). EdU incorporation was detected using a Click-iT® EdU kit (Thermo Fisher Scientific; #C10340). Immunofluorescence images were acquired using an Olympus FV1000 or a Leica SP8 scan confocal microscope. Image processing was performed with Fiji and Volocity software.

### RNA scope

RNAscope fluorescent in situ hybridizations (ISH) were conducted using the RNA-scope Multiplex Fluorescent Reagent Kit v.2 (Advanced Cell Diagnostics, 323100) following standard protocols provided by the manufacturer, using the following RNA-scope probes (ACDbio): Mm-Dot1L-C2 (#533431-C2); Mm-Hand1-C2 (#429651-C2); Mm-Irx4 (#504831)-

### Cardiomyocyte isolation

Embryonic and neonatal cardiomyocytes were isolated using a modified version of the a previously described protocol^38^. Briefly, embryonic or postnatal day 1 hearts were isolated and transferred into ice cold HBSS. Embryonic single cell suspensions from were obtained by performing eight rounds of enzymatic digestion (5 minutes each) with Collagenase type II (0.7mg/ml Worthington) at 37°C under agitation. Postnatal day 1 single cell suspensions were obtained by performing an overnight digestion with trypsin (0.5mg/ml) at 4°C followed by eight rounds of enzymatic digestion (5 minutes each) with Collagenase type II (1mg/ml Worthington) at 37°C. Cells were collected in cold medium containing fetal calf serum to stop the enzymatic reaction and centrifuged at 110 rcf to allow initial separation of cardiomyocytes from other cardiac cells. Cell preps were resuspended in 500μL of FACS buffer (HBSS, 5%FBS, 2.5mM EDTA) and 1μL of DAPI was added immediately prior to flow cytometry. Embryonic and postnatal day 1 live (DAPI negative) control and mutant cardiomyocytes were sorted based on the red fluorescence emitted by the Cre reporter tdTomato using an Influx Cell sorter (BD Biosciences) and collected in TRIZol reagent for RNA extraction or cross-linked as described below for chromatin analysis.

Postnatal day 10 cardiomyocytes were isolated using a Langerdoff system using Collagenase type II (0.7mg/ml Worthington). After perfusion cells were dissociated from the heart, collected in conical tubes and allowed to settle by gravity in order to obtain separation of viable, rod-shaped cardiomyocytes from dead cardiomyocytes and from interstitial cells of the heart. For histological analysis, isolated cardiomyocytes were fixed in 4%PFA and processed for immunostaining.

### Protein isolation and Western Blot analysis

Total protein extracts were prepared by lysing samples in RIPA buffer. Protein lysates in Laemmli buffer were separated by electrophoresis on 12% SDS-PAGE gels and transferred for 2 hours at 4°C on to a PVDF membrane (BioRad). After blocking for an hour in 5% dry milk, membranes were incubated overnight at 4°C with the primary antibody in blocking buffer. The following primary antibodies were used: H3 dimethyl Lys79 (abcam #ab3594) and anti-H3 (abcam #ab1791). Blots were washed and incubated with a horseradish peroxidase (HRP)-conjugated secondary antibody generated in Rabbit (1:2000; Cell Signaling Technology #7074) for 1.5 hours at room temperature. Immunoreactive protein bands were visualized using an enhanced chemiluminescence reagent (Thermo Fisher Scientific). Protein quantification was achieved using ImageJ software.

### Quantification of proliferation by Flow Cytometry

FACS quantification of rates of EdU incorporation was carried out using littermate Dot1L cKOs and Ctrls. EdU detection was done in cell suspensions using the Click-iT® EdU Alexa 647 kit (Thermo Fisher Scientific; C10340), according to the manufacturer’s instructions. TdTomato signal was used to discriminate cardiomyocytes from other lineages of cardiac cells. Stained cells were analyzed using a FACS Canto II flow cytometer (BD Bioscience). DIVA and FlowJo software (BD Pharmingen) were used for data acquisition and analysis.

### RNA extraction, qRT-PCR and RNA-seq

RNA was extracted using Tryzol (Invitrogen #15596026) and Direct-zol RNA Kits (Zymo Research #R2061) following instructions provided by the manufacturers. All transcriptomics analyses (qPCR or RNA-seq) were performed on FACS-sorted cardiomyocytes. For E16.5 and P1, biological replicates were prepared from single hearts. For E12.5, preparation of each biological replicate required pooling of hearts with same genotypes. For each stage analyzed, a minimum of 3 biological replicates, prepared from littermate Dot1L Ctrl and Dot1L cKO hearts, were used. For qRT-PCR experiments, cDNA was produced using the SuperScript VILO cDNA Synthesis Kit (Invitrogen #11754050). qRT-PCR was performed using SYBR Select Master Mix for CFX (Applied biosystems #4472952) on a Bio-Rad CFX96 Real-Time PCR system using the following primers: DOT1L-Ex2-Fw 5’-TGCTGCTCATGAGATTATTGAGA-3’ (primer hybridizing within the loxP-flanked exon2 of the floxed *Dot1l* allele); DOT1L-Ex4-Rev 5’- ATGGCCCGGTTGTATTTGTC-3’; 18s-Fw 5’-AAATCAGTTATGGTTCCTTTGGTC-3’; 18s- Rev 5’-GCTCTAGAATTACCACAGTTATCCAA-3’

For RNA-seq experiments libraries were generated from 25ng of RNA using the TruSeq Stranded mRNA library Prep kit (Illumina# 20020594) and sequenced on a HiSeq 4000 System (Illumina) using a single read 50 protocol.

### ChIP-seq

ChIP-seq was essentially performed as described^39^. Briefly, cells were fixed in in 1% formaldehyde/PBS for 10 minutes at room temperature, the reactions quenched by adding 2.625 M glycine to 125 mM final 20% BSA to 0.5% final and cells pelleted by centrifugation for 5 minutes at 1,000 g, 4°C. Cells were washed twice with ice-cold 0.5% BSA/PBS and cell pellets were snap-frozen in liquid nitrogen and stored at −80°C. Fixed cells were thawed on ice, resuspended in 500 µl ice-cold buffer L2 (0.5% Empigen BB, 1% SDS, 50 mM Tris/HCl pH 7.5, 1 mM EDTA, 1 × protease inhibitor cocktail) and chromatin was sheared to an average DNA size of 300–500 bp by administering 7 pulses of 10 s duration at 13 W power output with 30 s pause on wet ice using a Misonix 3000 sonicator. The lysate was diluted 2.5-fold with ice-cold L2 dilution buffer (20 mM Tris/HCl pH 7.4@20°C, 100 mM NaCl, 0.5% Triton X-100, 2 mM EDTA, 1 × protease inhibitor cocktail), and one percent of the lysate was kept as ChIP input. For each immunoprecipitation, aliquots of diluted lysate equivalent to 150,000 to 1 million cells, 20 µl of Dynabeads Protein A (for rabbit polyclonal antibodies) and 2 µg anti H3K79me2 antibody (Abcam #ab3594) were combined and rotated overnight at 8 RPM and 4°C. The following day, beads were collected on a magnet and washed three times each with wash buffer I (10 mM Tris/HCl pH 7.5, 150 mM NaCl, 1% Triton X-100, 0.1% SDS, 2 mM EDTA), wash buffer III (10 mM Tris/HCl pH 7.5, 250 mM LiCl, 1% IGEPAL CA-630, 0.7% Deoxycholate, 1 mM EDTA) and twice with ice-cold TET (10 mM Tris/HCl pH7.5, 1 mM EDTA, 0.2% Tween-20). Libraries were prepared directly on the antibody/chromatin-bound beads: after the last TET wash, beads were suspended in 25 µl TT (10 mM Tris/HCl pH7.5, 0.05% Tween-20), and libraries were prepared using NEBNext Ultra II reagents according to the manufacturer’s protocol but with reagent volumes reduced by half. DNA was eluted and crosslinks reversed by adding 4 µl 10% SDS, 4.5 µl 5 M NaCl, 3 µl EDTA, 1 µl proteinase K (20 mg/ml), 20 µl water, incubating for 1 h at 55°C, then overnight at 65°C. DNA was cleaned up by adding 2 µl SpeedBeads 3 EDAC in 61 µl of 20% PEG 8000/1.5 M NaCl, mixing and incubating for 10 minutes at room temperature. SpeedBeads were collected on a magnet, washed twice by adding 150 µl 80% EtOH for 30 s each, collecting beads and aspirating the supernatant. After air-drying the SpeedBeads, DNA was eluted in 25 µl TT and the DNA contained in the eluate was amplified for 12 cycles in 50 µl PCR reactions using NEBNext High-Fidelity 2X PCR Master Mix or NEBNext Ultra II PCR master mix and 0.5 µM each of primers Solexa 1GA and Solexa 1GB. Libraries were cleaned up as above by adding 36.5 µl 20% PEG 8000/2.5 M NaCl and 2 µl Speedbeads, two washes with 150 µl 80% EtOH for 30 s each, air-drying beads and eluting the DNA into 20 µl TT. ChIP library size distributions were estimated following 2% agarose/TBE gel electrophoresis of 2 µl library, and library DNA amounts measured using a Qubit HS dsDNA kit on a Qubit fluorometer. ChIP input material (1 percent of sheared DNA) was treated with RNase for 15 min at 37°C in EB buffer (10 mM Tris pH 8, 0.5% SDS, 5 mM EDTA, 280 mM NaCl), then digested with Proteinase K for 1 h at 55°C and crosslinks reversed at 65°C for 30 min to overnight. DNA was cleaned up with 2 µl SpeedBeads 3 EDAC in 61 µl of 20% PEG 8000/1.5 M NaCl and washed with 80% ethanol as described above. DNA was eluted from the magnetic beads with 25 µl of TT and library prep and amplification were performed as described for ChIP samples. Libraries were sequenced single-end for 75 cycles (SE75) to a depth of 20-25 million reads on an Illumina NextSeq 500 instrument.

### Bioinformatic analyses RNA-seq

Sequencing reads were processed to remove Illumina barcodes and aligned to the UCSC *Mus musculus* reference genome (build mm10) using STAR v.2.5.1b with default parameters^40^.

ReadsPerGene.out.tab files were then processed with edgeR^41^. RNA expression was calculated in reads per kilobase per million mapped reads (RPKM) considering the sum of exon length.

Differentially expressed coding genes were selected based on the following parameters: FDR ≤0.05, RPKM ≥1 in at least one biological condition, and log2 Fold Change ≤ −0.5 for genes downregulated and ≥ 0.5 for genes upregulated in Dot1L cKOs vs Dot1L Ctrls. Pathway analysis was performed using METASCAPE^42^.

### ChIP-seq alignment and peak-calling

All analyses were performed using the mouse reference genome GRCm38 (mm10) and the gencode gene annotation version vM25. Bowtie2^43^ was applied to align the fastq files to the mouse reference genome. First, the required index structure was built using: *bowtie2-build -f -- seed 123 --threads 20 Mus_musculus.GRCm38.dna.primary_assembly.fa mouse_GRCM38_mm10*. Since the reads where not paired, we ran for each fastq file: *bowtie2 -x mouse_GRCM38_mm10 -U <fastq-file> -S <output-file-name>.sam -q -t -p 30*. Conversion of the resulting files (sam) to bam format was done using samtools (version samtools 1.10)^44^. To allow easy visualization in a genome browser, bam files were converted to bigWig (bw) files using deeptools bamCoverage function. Next peak-calling was performed with MACS2 (version macs2 2.2.7.1)^45^: *macs2 callpeak -t <treatment>.bam -c <input>.bam -n <prefix-name> -- outdir <output-dir> -f BAM -g 1.87e9 -B*, where <treatment> is either the aligned reads of the Dot1L Ctrl or Dot1L cKO and <input> the corresponding input signal. For all following analyses we used the narrowPeak files.

To compute differential H3K79me2 peaks between Ctrl and Dot1L cKOs, we applied DiffBind (version 2.10.0 bioconductor-diffbind)^46^ using the resulting narrow peak files from the peak calling of Ctrl and cKO cardiomyocytes (both replicates) and the corresponding bam files of the H3K79me2 ChIP-seq and input.

### Determining genomic distribution of H3K79me2 peaks

To determine the number of H3K79me2 peaks overlapping with distinct genomic regions, the gene annotation file was split into exon, UTR and gene regions. The most 5’ transcription start site (TSS) for all mouse genes was downloaded from biomart^47^ and a promoter region of length 400 bp, centered around the most 5’ TSS, was determined. The number of H3K79me2 peaks overlapping either with exon, UTR or promoter regions was computed using the intersect function (used flags -f 0.1 -F 0.1 -e) from bedtools (v2.25.0)^48^. Subsequently, based on this information, peaks overlapping with introns and intergenic regions were deduced. Peaks were assigned as introns when they overlapped a gene but not an exon. Peaks that did not intersect with any of the aforementioned regions (exons, UTRs, promoters and introns) were classified as intergenic. Results from these analyses were plotted in the diagrams shown in Figure 3b (corresponding to E16.5) and Supplementary Figure 3b (corresponding to P1).

### Computing coverage and fraction of the gene body covered by H3K79me2

The profiles of the H3K79me2 signal at the gene body were computed using deeptools^49^. We used the .bam files resulting from the bowtie2 analysis and the up- and downregulated genes in bed-file format (with strand information).

First, we used deeptools bamCompare to determine the mean signal of the two replicates of Ctrl and cKO for each time point: *bamCompare -b1 <treatment_rep1>.bam -b2 <treatment_rep2>.bam -o <treatment>_mean_rep1_rep2.bw -of bigwig --scaleFactorsMethod None --operation mean --effectiveGenomeSize 2652783500 -p 25 --normalizeUsing RPKM -- binSize 20 --skipNonCoveredRegions*, where <treatment> is either Ctrl or cKO.

Next, we visualized the data using deeptools computeMatrix and plotProfile functionalities: *computeMatrix scale-regions -S Ctrl_mean_rep1_rep2.bw cKO_mean_rep1_rep2.bw -R downregulated_genes_FDR_0.05.bed upregulated_genes_FDR_0.05.bed -o inputProfile.mat.gz --endLabel TTS --beforeRegionStartLength 2000 --afterRegionStartLength 2000 --regionBodyLength 5000 -p 20 --skipZeros*

*plotProfile -m inputProfile.mat.gz -out profile.pdf --perGroup --startLabel TSS --endLabel TTS - -plotTitle “Average distribution of H3K79me2 relative to the distance from TSS and TTS” -- samplesLabel “Ctrl (upregulated genes)” “cKO (upregulated)” --regionsLabel “downregulated genes” “upregulated genes” --plotFileFormat pdf*

To compute the fraction of the gene body which is covered by H3K79m2, we applied bedtools (bedtools v2.25.0) coverage function. Given a set of genomic regions and a .bam file, the function computes per genomic region the number of reads that overlap and the fraction of bases that have a non-zero coverage based on the .bam file. We determined the number of overlapping reads and the fraction for all annotated mouse genes for Ctrl and cKO. Next, the mean fraction (of rep1 and rep2) of the up- and downregulated genes was computed. The density plots are based on all up- and downregulated genes.

### Calling regulatory interactions with H3K27ac ChIP-seq and Hi-C and differential H3K79me2 enhancer analysis

For the identification of enhancer regions H3K27ac ChIP-seq data from C57BL/6 heart tissue was downloaded from ENCODE for both timepoints E16.5 (ENCFF153DSU) and P1 (ENCFF463XJT). A Hi-C matrix of mouse cardiomyocytes was downloaded from GEO (GSM2544836)^30^. The Hi-C matrix was normalized using Knight-Ruiz normalization with the Juicebox dump command (version 1.22.01)^50^ with a resolution of 5000 bp for each chromosome. *java -jar juicer_tools_1.22.01.jar dump observed KR <hic-file> chr$ chr$ BP 5000 <output-file-name>*

The regulatory interactions between promoter and candidate enhancers were assessed with a self-implemented version of the ABC-score of Fulco et al.^26^. For all genes on the autosomes (GRCm38p6) the most 5’ TSS was taken and all candidate enhancers in a 5Mb window centered on the TSS were scored according to the following equation:

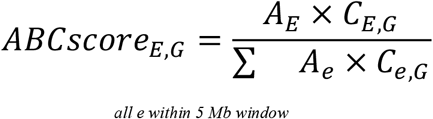

The interaction of an enhancer (*E*) with a gene (*G*) is described by the activity of the enhancer (*AE*) and the contact to the gene (*CE,G*). The ABC-score returns the relative contribution of an interaction in relation to all other interactions of that gene. The enhancer activity is the signal of the H3K27ac ChIP-seq peak and is scaled so that the maximum value for all enhancers within a gene window is 100. The contact (*CE,G*) between an enhancer and a gene’s TSS is the contact frequency of the respective bins in the normalized Hi-C matrix and is also scaled to a maximum of 100 for all enhancers within the window. All scored candidate enhancer-gene interactions were ranked according to the ABC-score and the top 5% were used for further analyses.

Chromosomes X and Y where excluded from the analysis. An enhancer was considered to be differential for H3K79me2 if ≥ 10% of the enhancer’s length was covered by a differential H3K79me2 peak. Statistical differences between the cumulative log2(fold-change of gene expression) distributions were calculated pairwise with a two-tailed Kolmogorov-Smirnov test (SciPy 1.4.1, stats module, ks_2samp function).

### Motif enrichment analysis

To identify transcription factors likely regulating the activity of K79-REs associated to genes differentially expressed in Dot1L cKOs, a motif enrichment analysis was performed. To this end, REs overlapping with a H3K79me2 peak and associated to differentially expressed genes were divided into two groups: those associated with genes downregulated in Dot1L cKOs (activating K79-REs, N=1355) and those associated with genes upregulated in Dot1L cKOs (silencing K79-REs, N=1047). TF binding motifs (total of 515) were downloaded from the JASPAR database^51^ and only those significantly expressed (RPKM ≥ 1 in our RNA-seq dataset) in E16.5 cardiomyocytes (a total of 367 TFs) were considered in subsequent analyses. Using TRAP^52^, a TF-affinity value per TF for each RE sequence was computed. The value is defined as the sum over all binding site probabilities of the given TF for the current sequence. The TRAP analysis was performed separately for the RE sequences of the up- and downregulated genes. A one-tailed Mann-Whitney test (using R’s Wilcox test function, confidence level of 0.975) to identify TFs enriched in the K79-REs of upregulated genes versus K79-REs of downregulated genes and vice versa was performed. The resulting p-values were adjusted by applying the Benjamini– Hochberg procedure. All TFs with an adjusted p-value of 0.05 were considered significant.

### Quantification Experiments and Statistical Analysis

Statistical significance of differences in the survival of *Dot1L* mice was assessed using Kaplan-Meier survival analysis with the log-rank method of statistics. In all other graphs, data are expressed as mean ± SEM, with a minimum of 3 biological replicates (the exact replicate number is described in the legend to each figure). Statistical significance of differences among groups was tested by 2-tailed Student’s t-test. A value of P ≤0.05 was considered statistically significant. * P≥0.05, **P≥0.01. Analyses were performed with GraphPad Prism software.

## Supporting information

Supplementary File 1_E16+P1_RNA-seq

Supplementary File 2_E16+P1_DiffBind

Supplementary File 3_E16+P1_REs and ABC analysis

Supplementary File 4_E16+P1_REs associated with gene expression

## Nonstandard Abbreviations and Acronyms

DOT1L: disruptor of telomeric silencing 1
H3K79me: methylation of Lysine 79 of Histone 3
H3K27ac: acetylation of Lysine 27 of Histone 3

## Data Availability

All RNA-seq and ChIP-seq data that support the finding of this study have been deposited in the Gene Expression Omnibus (GEO) under accession number code GSE184192. Previously published ChIP-seq and Hi-C data that were re-analyzed here are available under accession codes ENCFF153DSU and ENCFF463XJT from ENCODE and GSM2544836 from GEO. Source data are provided with this paper. All other data supporting the findings of this study are available from the corresponding authors upon reasonable request.

## Sources of Funding

PC was supported by the Marie Curie International Outgoing Fellowship within the 7th European Community Framework Program under grant agreement No 623739 - The Cardiac Code and a grant from the German Center for Cardiovascular Research (DZHK). NG-C was supported by a junior research group grant from the German Center for Cardiovascular Research (DZHK). JC and SME are funded by grants from the National Heart, Lung, and Blood Institute, and the Leducq Foundation.

## Acknowledgments

Confocal microscopy was performed at the University of California, San Diego School of Medicine Microscopy Core Facility, which is supported by the grant P30 NS047101. Tim Mohun from the Francis Crick Institute of London (UK) kindly provided the xMlc2-Cre mice. Ian James Jackson from the Institute of Genetic and Molecular Medicine, University of Edinburgh (UK) kindly provided the Rosa26-Fucci2a mice.

## Disclosures

None to declare.

## Contributions

P.C. and N.G.-C. conceived the study and performed most of the experiments, analyzed data, acquired funding sources and prepared the manuscript. S.P., V.L, L.Z., R.C. and S.S. contributed to experiments. M.H, G.F. and S.H. performed ChIP-seq experiments. P.K analyzed RNA-seq datasets. N.B and D.H. under supervision by M.H.S. performed additional bioinformatics analyses on ChIP-seq. J.C and G.C. contributed to project supervision. S.M.E. conceived and supervised the study and acquired funding sources. All authors commented and edited the manuscript.

## Notes

### Competing Interest Statement

The authors have declared no competing interest.

