## Supplementary Figures for "The epigenetic modifier DOT1L regulates gene regulatory networks necessary for cardiac patterning and cardiomyocyte cell cycle withdrawal"

**Supplementary Figure 1 - Cattaneo et al.**

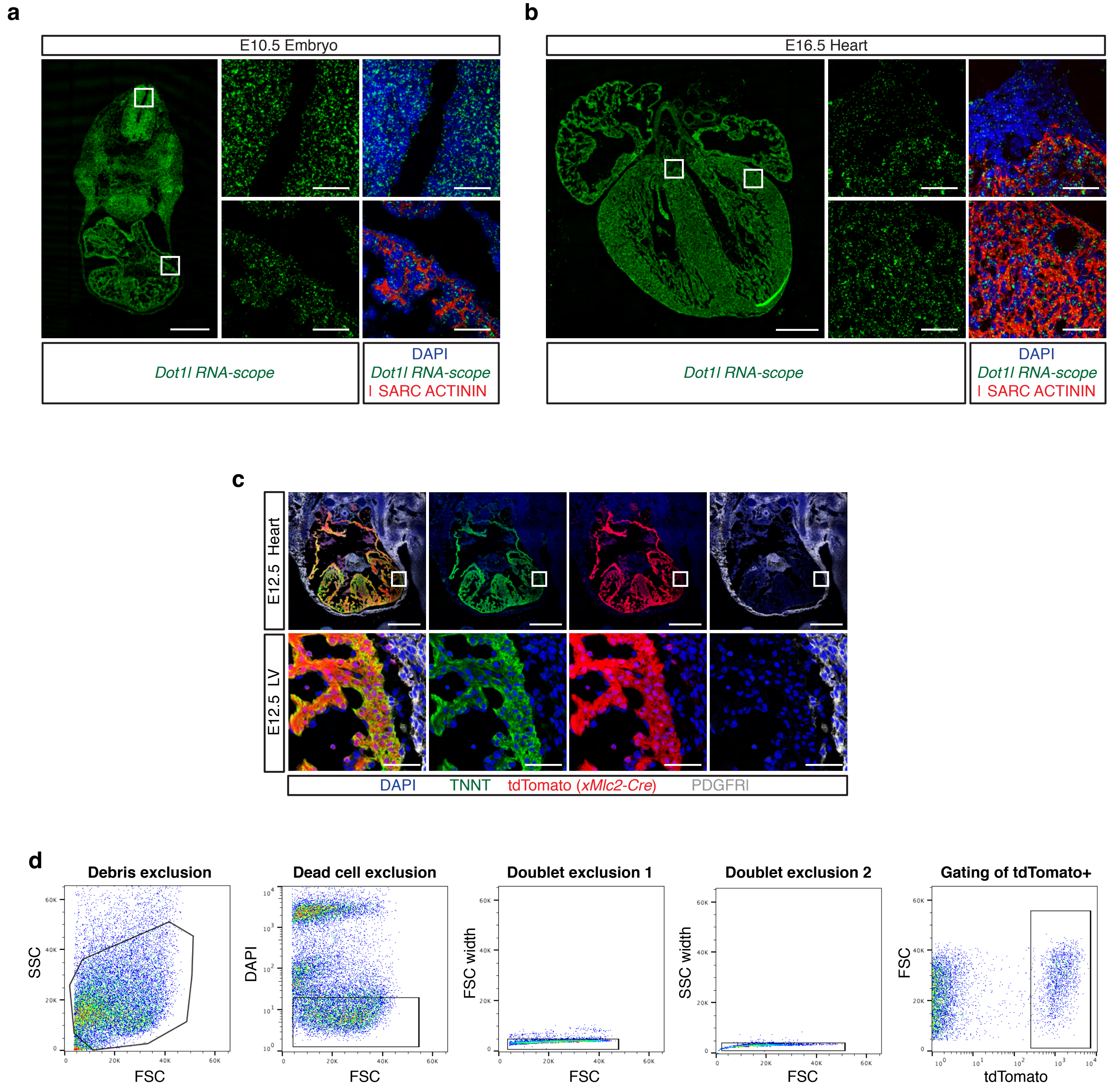


**e**

| Stage | N litters | N mice | Without Cre *Dot1l*$^{fl/+}$ | Without Cre *Dot1l*$^{S/fl}$ | *xMlc2-Cre* *Dot1l*$^{fl/+}$ | *xMlc2-Cre* *Dot1l*$^{S/fl}$ |
|---|---|---|---|---|---|---|
| E12.5 | 23 | 190 | 30.0% | 24.2% | 21.6% | 24.2% |
| E14.5 | 8 | 65 | 27.7% | 13.8% | 30.8% | 27.7% |
| E16.5 | 30 | 255 | 24.7% | 25.5% | 25.9% | 23.9% |
| E18.5 | 2 | 19 | 15.8% | 10.5% | 47.4% | 26.3% |
| P0 | 33 | 261 | 25.3% | 26.4% | 27.2% | 21.1% |

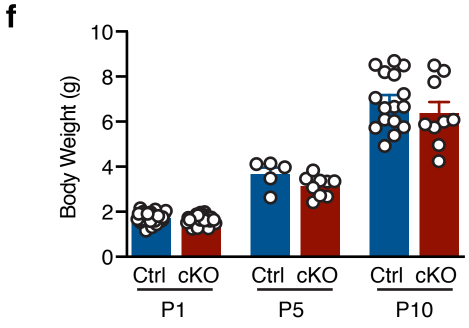

**Supplementary Figure 1, related to Figure 1: Patterns of *Dot1l* expression and strategy for its conditional deletion in cardiomyocytes from early embryonic timepoints. a-b)** RNA-scope detection of *Dot1l* transcripts in distinct embryonic stages. *Dot1l* was ubiquitously expressed in E10.5 embryos (a). In the developing E16.5 heart, *Dot1l* mRNA could be detected in all cell types, but cardiomyocytes (labeled by αSARCOMERIC ACTININ in red) displayed higher abundance of *Dot1l* transcripts than non-myocyte lineages (b) Scale bar = 500μm for low magnification panels and 50μm for high magnification of boxed areas). **c)** The *xMlc2-Cre* allele was highly specific and efficient in promoting recombination in embryonic cardiomyocytes, as evidenced from the extensive overlap between the tdTomato recombination reporter signal (Cre+ cells, in red) and the cardiomyocyte marker Troponin T (TNNT, in green), as well as the lack of colocalization between tdTomato and the marker of mesenchymal lineages PDGFRα (white). To facilitate visualization of patterns, in addition to a 4-colour merge (left panels), images are also provided for each individual protein. (scale bar = 500 μm low magnification top panels; 50 μm for high magnification of boxed areas). **d)** FACS strategy to isolate highly pure populations of cardiomyocytes based on the signal emitted by the red fluorescent protein tdTomato (expressed only in cells hit by the *xMlc2-Cre*). **e)** Observed genotype distribution of live embryos recovered from mattings between *XMlc2-Cre;Dot1lΔ/+* and *Dot1lfl/fl* mice. No significant deviations from the expected Mendelian distribution (25% for each genotype) were observed until birth. **f)** No significant body weight differences were observed between control and cKO mice at P1, P5 and P10.

**Supplementary Figure 2 - Cattaneo et al.**

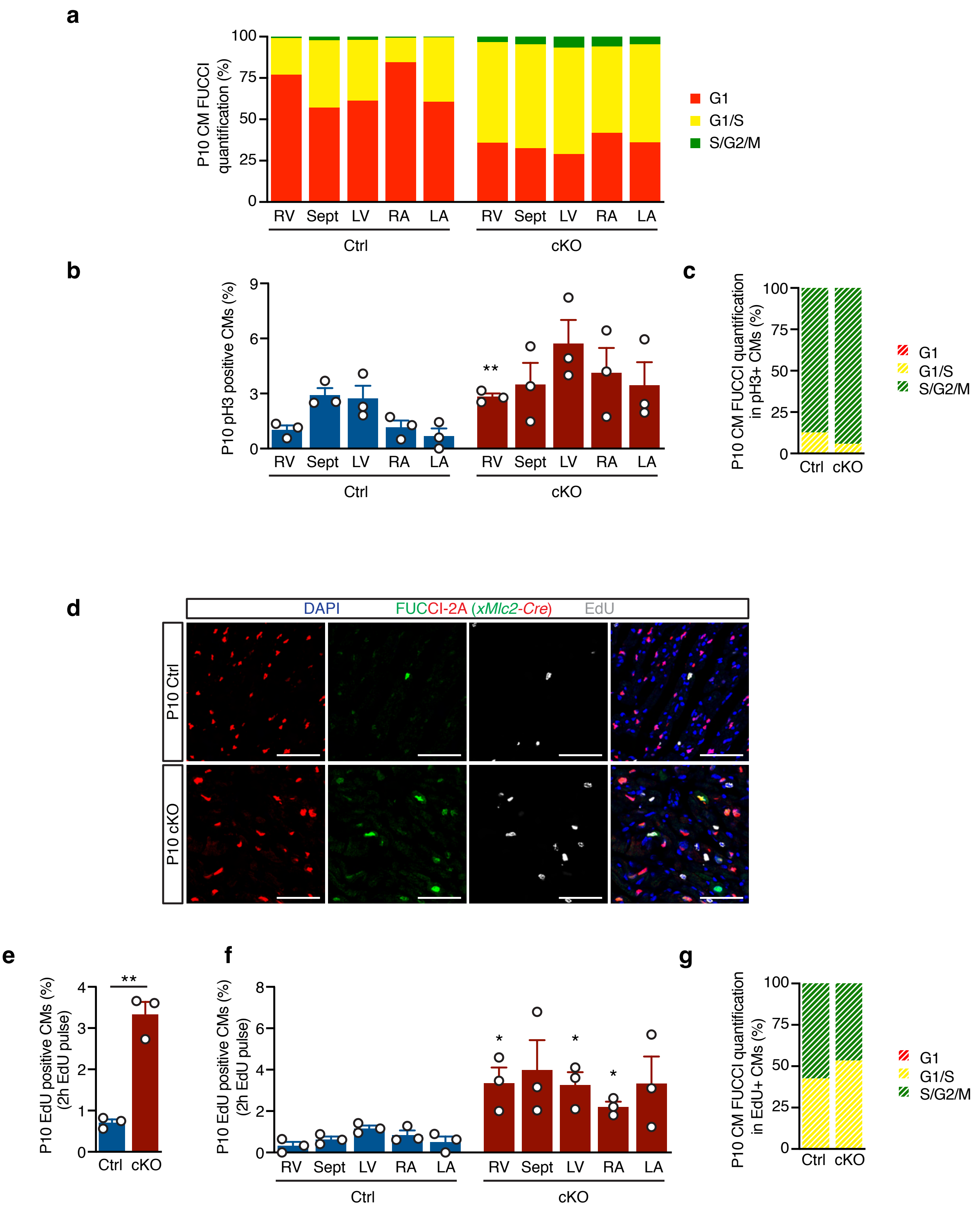

**Supplementary Figure 2, related to Figure 5: Dot1L cKO cardiomyocytes fail to undergo neonatal cell cycle withdrawal. a)** Quantification of cardiomyocyte cell cycle phase distribution across the distinct cardiac chambers in histological sections from P10 Dot1L Ctrl and cKO hearts on a *Rosa26-Fucci2A* background (Cre-dependent cell cycle indicator). Red only signal represents cardiomyocytes in G1; red+green (yellow) signal represents cardiomyocytes in G1/S; green only signal represents cardiomyocytes in S/G2/M. Right ventricle (RV), septum (Sept), left ventricle (LV), right atrium (RA) and left atrium (LA) (N=3 per compartment). Increased ratios of proliferative cardiomyocytes (G1/S and S/G2/M) were found in all compartments of the Dot1L cKO heart. **b)** Quantification of mitotic cardiomyocytes (pH3+) in histological sections of P10 Dot1L Ctrl and cKO hearts (*Rosa26-Fucci2A* background). Compartment-specific data are presented as percentage of pH3+ cardiomyocytes over total cardiomyocytes (N=3; mean± SEM). Increased percentage of pH3+ cardiomyocytes were found in all compartments of the Dot1L cKO heart. **c)** Distribution of pH3+ cells according to the cell cycle stage indicated by the FUCCI2A reporter. In both genotype groups the majority of pH3+ cardiomyocytes corresponded to cardiomyocytes in S/G2/M (green only). **d-e-f)** Immunofluorescence images (d) and corresponding quantification (e,f) showing signals of EdU incorporation (white) on histological sections from P10 Dot1L Ctrl and cKO hearts on a *Rosa26-Fucci2A* indicator background (Scale bars = 50μm). At P10, when analyzing all cardiac chambers together (e), or distinct compartments separately (f), Dot1L cKO hearts displayed significantly higher percentages of EdU+ cardiomyocytes than their Ctrl counterparts (mean of 1022 CMs counted per heart; N=3; **P≤0.01; mean± SEM). **g)** Distribution of EdU+ cells according to the cell cycle stage indicated by the FUCCI2A reporter. In both genotype groups all EdU+ cardiomyocytes corresponded to cardiomyocytes in G1/S (yellow) or S/G2/M (green only).

**Supplementary Figure 3 - Cattaneo et al.**

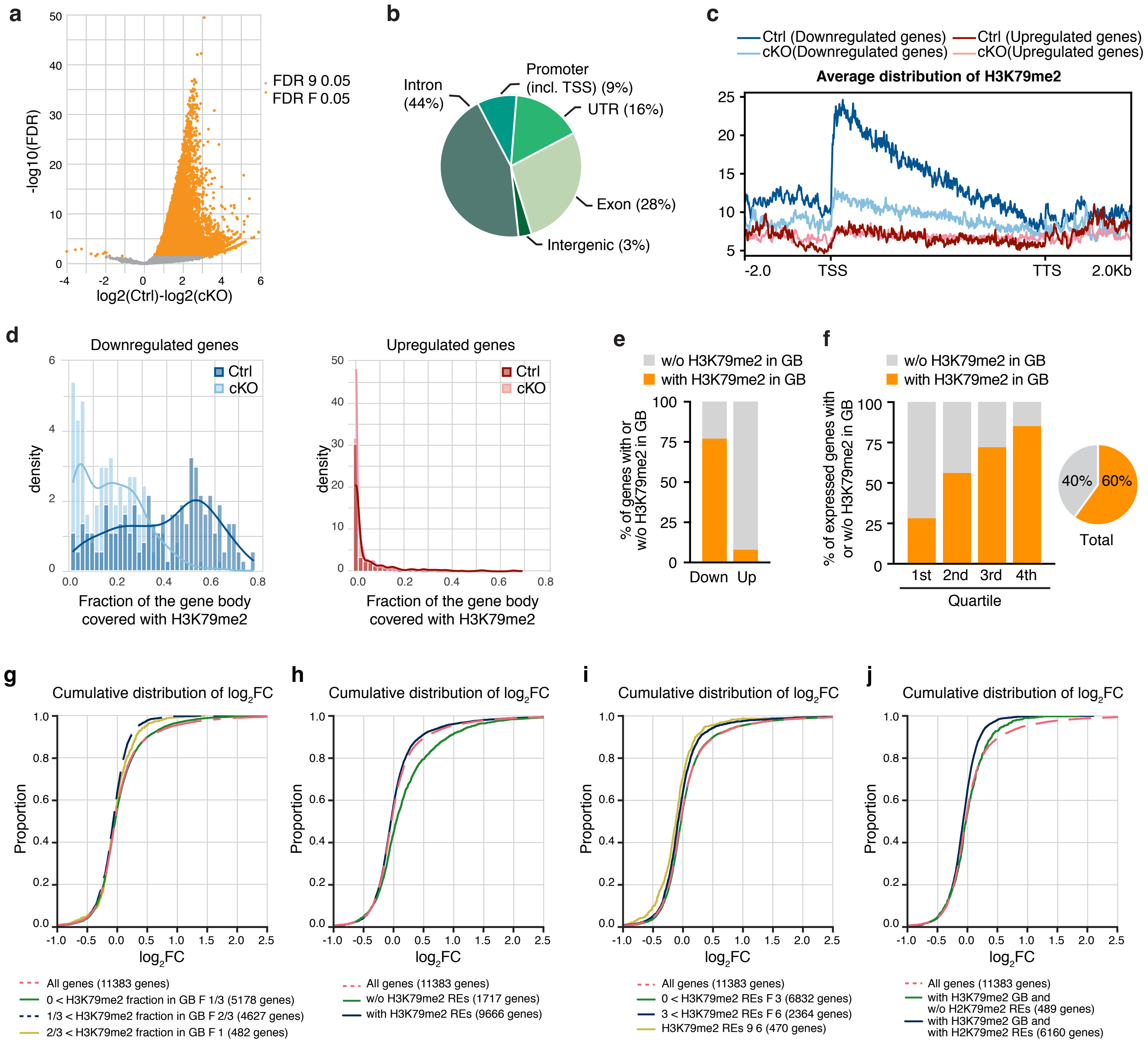

**Supplementary Figure 3, related to Figure 5: Mechanisms of gene expression regulation by DOT1L/H3K79me2 in postnatal day 1 cardiomyocytes. a)** Volcano plot displaying H3K79me2 ChIP-seq peaks significantly enriched in P1 Ctrl vs cKO cardiomyocytes. **b)** Pie chart indicating the genomic distribution of differential H3K79me2 ChIP-seq peaks in P1 cardiomyocytes. **c)** Metagene profiles showing the average distribution of H3K79me2 input-normalized tag density relative to Transcription Start Site (TSS) and Transcription Termination Site (TTS) with ±2Kb flanking regions. Genes downregulated in P1 Dot1L cKOs had, in Dot1L Ctrl cardiomyocytes, high levels of H3K79me2 in the vicinity of the TSS that progressively decreased towards the TTS. Genes upregulated in P1 Dot1L cKOs did not show significant level of H3K79me2 in Ctrls nor cKOs. **d)** Fraction of gene body covered with H3K79me2 in downregulated genes (left graph) and in upregulated genes (right graph) indicating that most genes downregulated in P1 Dot1L cKOs had, in Ctrl cardiomyocytes, H3K79me2 covering the majority of the gene body, whereas the majority of genes upregulated in cKOs didn't have H3K79me2 gene body coverage in Ctrl cardiomyocytes. **e)** Graph representing the percentage of down- and up-regulated genes in P1 cKO cardiomyocytes with (Coverage ≥ 50 reads and Fraction of gene body ≥ 0.2) or without (Coverage < 50 reads and Fraction of gene body < 0.2) gene body H3K79me2 in P1 control cardiomyocytes. **f)** Percentage of genes with H3K79me2 in the gene body (GB) (Coverage ≥ 50 reads and Fraction of gene body≥ 0.2) or without H3K79me2 in the gene body (Coverage <50 reads and Fraction of gene body < 0.2). This modification was abundant amongst highly expressed genes ($4^{th}$ quartile of RNA expression) and progressively decreased towards the lower quartiles of expression. Globally more than half (60%) of all genes expressed in P1 cardiomyocytes had gene body H3K79me2. **g-h-i-j)** Cumulative log2FC distribution analyses in different categories of genes in P1 cardiomyocytes: genes with different levels of H3K79me2 fraction in the gene body (GB) (g); genes with and without H3K79me2/H3K27ac positive regulatory elements (REs) (h); genes with increasing numbers of H3K79me2/H3K27ac positive REs (i); genes with a combination or none of H3K79me2 in GB and REs (j). DOT1L regulated expression of target genes via H3K79me2 in GB and REs and genes associated with both types of H3K79me2 (GB and RE) were more downregulated in cKOs than those containing exclusively gene body H3K79me2. A Kolmogorov-Smirnov test was applied to assess statistical significance of differences between distribution of gene groups.

**SUPPLEMENTARY FILES:**

**Supplementary File 1: Integrated transcriptomics and epigenomics analyses in E16.5 and P1 cardiomyocites.**

**a,b)** RNA-seq and H3K79me2 ChIP-seq results comparing the transcriptomes and epigenomes of cardiomyocytes FACS sorted from E16.5 (a) or P1 (b) Dot1L cKO vs Ctrl hearts.

Column headlines and description: *Gene ID, Ensembl ID, UCSC ID* unique identifiers of analyzed genes; *biotype* = category of analyzed genes; *chr, start, end* = coordinates of analyzed genes; *DEG* (Differential Expressed Genes), *logFC* (log Fold Change), *log CPM* (log Count Per Million), *LR* (Likelihood Ratio) , *PValue, FDR* (False Discovery Rate) refers to the edgeR differential RNA-seq analysis (DOWN = significantly downregulated genes in cKO vs Ctrl with log2FC ≤ 0.5 and FDR ≤ 0.05, UP = significantly upregulated genes in cKO vs Ctrl with log2FC ≥ 0.5; FDR ≤ 0.05, NS = not significantly differentially expressed genes); *Ctrl Coverage, cKO Coverage* = H3K79me2 reads in the gene body in Ctrls and cKOs (mean of replicates); *Ctrl Fraction, cKO Fraction* = fraction of gene body that is covered by H3K79me2 reads in Ctrls and cKOs (corrected for input, mean of replicates).

**Supplementary File 2: H3K79me2 ChIP-seq analyses.**

**a,b)** DiffBind analyses determining differential H3K79me2 ChIP-seq peaks in E16.5 (a) and P1 (b) Dot1L cKO vs Ctrl sorted cardiomyocytes. Differential bound sites between Ctrl and cKO with FDR≤ 0.05 are shown, sorted for fold enrichment.

Column headlines and description: *Width* = Length of site; *Conc* = Mean read concentration over all the samples (the default calculation uses log2 normalized ChIP read counts with input read counts subtracted); *Conc_Ctrl* = Mean concentration over control group; *Conc_cKO* = Mean concentration over cKO group; *Fold* = difference in mean concentrations between Dot1L Ctrl and Dot1L cKO groups; *p.value* = P-value confidence measure for the identification of individual peaks as differentially bound; *FDR* = False Discovery Rate.

**Supplementary File 3: Genomic interactions underlying H3K79me2-dependent gene regulation. a,b)** Top 5% most relevant regulatory element-to-target gene interactions identified by the Activity-by-contact (AbC) model in E16.5 (a) and P1 (b) cardiomyocytes.

Column headlines and description: *Peak_Chr, Peak_Start, Peak_End*, refer to the coordinates of each H3K27ac ChIP-seq peak with significant interactions with target genes; *Interacting geneID* = gene interacting with the peak; *norm_signalValue* = H3K27ac peak signal normalized for length; *peakID* = internal ID for identification of H3K27ac peaks; *K79meDiff* = whether or not the H3K27ac peak overlaps with a differential H3K79me2 peak (no=0, yes=1); *K79meFC* = logFoldChange (Dot1L Ctrl versus Dot1L cKO) of the overlapping differential H3K79me2 peak(s) (from DiffBind analysis, Supplementary File 2). If more than one differential H3K79me2 peak overlaps with the H3K27ac peak, the log fold change for each of the H3K79me2 peaks is separated by an "|"; *scaledActivity*= Activity scaled so that the maximum of all peaks 5Mb around the gene is 100; *scaledContact* = Contact scaled so that the maximum of all peaks 5Mb around the gene is 100; *ABC-Score* = score of peak-gene interaction; *TSS-dist* = distance between the peak and the transcriptional start site (TSS) of its target gene; *ovPromoter* = if the H3K27ac peak overlaps with a promoter of a known gene, the ID of such gene(s) is listed in this column; *Location* = genomic location of the peak. In this classification "1.0_intergenic" means the peak does not overlap with any annotated gene. "1.0_genebody" means the peak is 100% within a gene. In this case, additional information is provided as to peak distribution across specific gene domains (promoter/exon/UTR).

**Supplementary File 4: Number of REs associated to gene expression. a)** Analysis in E16.5 cardiomyocytes **b)** Analysis in P1 cardiomyocytes.

Column headlines and description: *geneID, GeneName, chr,* and *TSS* (Transcription Start Site) refer to each gene expressed in FACS-sorted cardiomyocytes (E16.5, or P1, depending on the tab) and having at least one interaction within the top 5% regulatory element-to-target gene interactions identified by the AbC analysis; *DEG, logFC, FDR* refers to the RNA-seq analysis (DOWN = significantly downregulated in cKO vs Ctrl with log2FC ≤ - 0.5 and FDR ≤ 0.05, UP = significantly upregulated in cKO vs Ctrl with log2FC ≥ 0.5; FDR ≤ 0.05, NS = not significantly differentially expressed); *Ctrl Coverage, cKO Coverage* = H3K79me2 reads in the gene body in Ctrls and cKOs (mean of replicates); *Ctrl Fraction, cKO Fraction* = fraction of gene body that is covered by H3K79me2 reads in Ctrls and cKOs corrected for input (mean of replicates); *#All-K27-REs* = number of all associated regulatory elements; *#All-K27/K79-REs* = number of associated regulatory elements that overlap a differential peak for H3K79me2; *#Intragenic-K27-REs* = number of regulatory elements that are located intragenic; *#Intragenic-K27/K79-REs* = number of intragenic regulatory elements that overlap a differential peak for H3K79me2; *#Intergenic-K27-REs* = number of regulatory elements that are located intergenic; *#Intergenic-K27/K79-REs* = number of intergenic regulatory elements that overlap a differential peak for H3K79me2).
